# Conjugative transfer inhibition of IncA and IncC plasmids by pervasive SGI1-like elements via relaxosome assembly interference

**DOI:** 10.1101/2025.04.21.649884

**Authors:** Florence Deschênes, Romain Durand, Kevin T. Huguet, Nicolas Rivard, Chloé Chagnon, Doriane Planez, Vincent Burrus

## Abstract

Broad-host-range IncA and IncC (A/C) conjugative plasmids propagate multidrug resistance in bacteria and mobilize chromosomal resistance islands, including *Salmonella* Genomic Island 1 (SGI1) across genera. A/C plasmids usually mobilize SGI1 at very high frequencies while being inhibited by SGI1. Here, we identified a broadly conserved fertility inhibitor, which we named CtiC (for ’Conjugative Transfer Inhibition of IncC’), encoded by SGI1 and kin, that hampers A/C plasmid transfer. *ctiC* expression is both constitutive and activated during conjugative transfer of A/C plasmids. Our results indicate that the suppression of *ctiC* significantly enhances plasmid transfer, suggesting it counters a general improvement in conjugation mediated by the two genes *traHG*, located upstream of *ctiC* in SGI1. CtiC specifically prevents the cotransfer of the helper plasmid, which destabilizes SGI1 in transconjugants. Structural predictions revealed that CtiC resembles the C-terminus of the MOB_H12_-family relaxase TraI of A/C plasmids. Bacterial two-hybrid assays and relaxase domain substitutions show that CtiC interferes with relaxosome assembly by binding to the plasmid-encoded mobilization factor MobI, which recognizes the origin of transfer, thus preventing its interaction with TraI. Our work reveals a fertility inhibition mechanism that prevents relaxosome assembly and uncovers a functional domain in MOB_H12_ relaxases.

## Introduction

Bacterial genomes can host a rich and diverse bestiary of mobile genetic elements (MGEs). Self-transmissible MGEs, such as conjugative plasmids and integrative and conjugative elements (ICEs), facilitate the exchange of antibiotic-resistance genes (ARGs) between bacteria by spreading from host to host via conjugation, a mechanism relying on a type IV secretion system (T4SS) that injects their DNA into recipient cells (1–4). Other MGEs, called integrative and mobilizable elements (IMEs), lack the genes required for self-transmissibility and coopt the T4SS encoded by helper conjugative MGEs to propagate (5, 6).

Conjugative plasmids of the incompatibility group C (IncC) contribute to the pervasive spread of ARGs across taxonomic barriers in pathogenic species of *Enterobacteriaceae*, *Vibrionaceae*, *Yersiniaceae*, and *Morganellaceae* in healthcare settings, animal husbandry and the environment (7). IncC and related IncA plasmids (collectively referred to as A/C plasmids) share a ∼128-kb set of core genes (>93% identity) disrupted by insertions of variable DNA and ARGs (8–12). Among the conserved genes are those involved in plasmid replication and partition, conjugative transfer (*tra* genes and operons, and *mobI*), entry and surface exclusion, DNA repair, and regulation (*acaB*, *acaDC*, *acr1*, *acr2*) (8, 13–21). The conjugative transfer of A/C plasmids is initiated at the origin of transfer (*oriT*), a cis-acting locus located upstream of *mobI* (8, 19). The mobilization factor MobI is thought to bind to *oriT* and recruit the MOB_H12_ family relaxase TraI, which initiates the intercellular rolling-circle replication of the DNA to transfer.

Besides self-propagation, A/C plasmids promote the transmission of multiple IMEs, including the ARGs-bearing 43-kb *Salmonella* Genomic Island 1 (SGI1) (22–24). SGI1 and its variants typically integrate at the 3’ end of *trmE* in a broad range of bacterial species, including *Proteus mirabilis* (PGI1), *Acinetobacter baumannii* (AGI1), and *Vibrio cholerae* (GI-15) (25). Distant relatives of SGI1 have also been found integrated at alternative sites (*dusA* and *yicC*) (26). One of their representatives, IME*Vch*USA3 of *V. cholerae*, lacks ARGs but remains mobilizable by A/C plasmids. Despite encoding different integrases targeting different integration sites, all SGI1-related IMEs share a small set of conserved genes that play a crucial role in their mobilization. Among those, *mpsAB* encodes the relaxase that recognizes and processes their *oriT* independently of the relaxase TraI of A/C plasmids (27). Also conserved are *traG* (and often abutting gene *traH*) and *traN*, which encode distant orthologues of the corresponding mating pair formation protein TraG and mating pair stabilization protein TraN of A/C plasmids (13, 26). Past reports showed that SGI1 or its variants can drastically inhibit (up to 2- to 3-log decrease) the transfer of A/C plasmids and that transconjugants carrying both the IME and its helper plasmid are rare (14, 22, 28–30). Competition for the T4SS is an attractive hypothesis that could explain part of the inhibition of helper plasmid transfer by an IME. However, competition alone is unlikely to result in such a drastic effect unless SGI1-like IMEs encode factors that skew the T4SS to selectively enhance IME mobilization or inhibit helper plasmid transfer, a phenomenon evocative of fertility inhibition observed between unrelated plasmids (31).

Here, we leveraged transposon-directed insertion sequencing (TraDIS) to identify SGI1 genes that negatively affect the transfer of a helper IncC plasmid. We report that a small, conserved gene located immediately downstream of *traG* is solely responsible for inhibiting the transfer of the helper A/C plasmids. Structural modelling and bacterial double-hybrid assays suggest fertility inhibition results from relaxosome assembly interference. Furthermore, our results indicate that fertility inhibition offsets a general improvement of the IncC conjugative apparatus promoted by SGI1, likely to prevent the cotransfer of the IME and its helper plasmid, which otherwise destabilizes the IME in transconjugants.

## Materials and methods

### Bacterial strains, plasmids, and media

The bacterial strains and plasmids used in this study are described in Table 1. Bacterial strains were grown at 37°C on Luria-Bertani (LB) agar or with agitation in LB broth. Bacterial strains were maintained at −75°C in LB broth containing 20% (vol/vol) glycerol. Antibiotics were used at the following concentrations: ampicillin (Ap), 100 μg/ml; kanamycin (Km), 50 μg/ml; spectinomycin (Sp), 50 μg/ml; rifampicin (Rf), 50 μg/ml; chloramphenicol (Cm), 20 μg/ml; nalidixic acid (Nx), 40 μg/ml; tetracycline (Tc), 12 μg/ml. For induction of gene expression, bacterial cultures were supplemented with 0.02% (wt/vol) L-arabinose or 1 mM isopropyl β-D-1-thiogalactopyranoside (IPTG). When necessary, LB agar was supplemented with 40 μg/ml 5-bromo-4-chloro-3-indolyl β-D-galactopyranoside (X-gal).

**Table 1.**
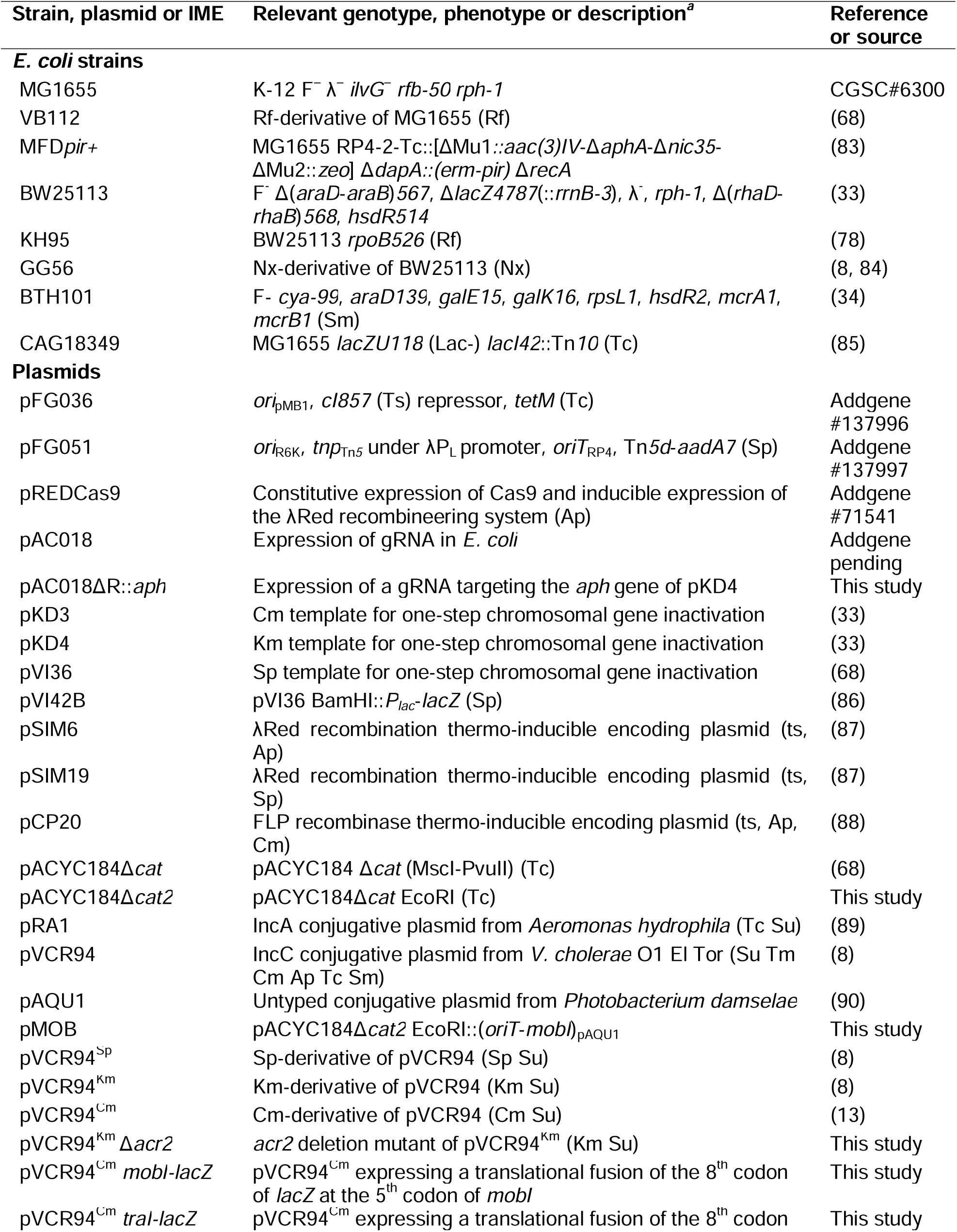
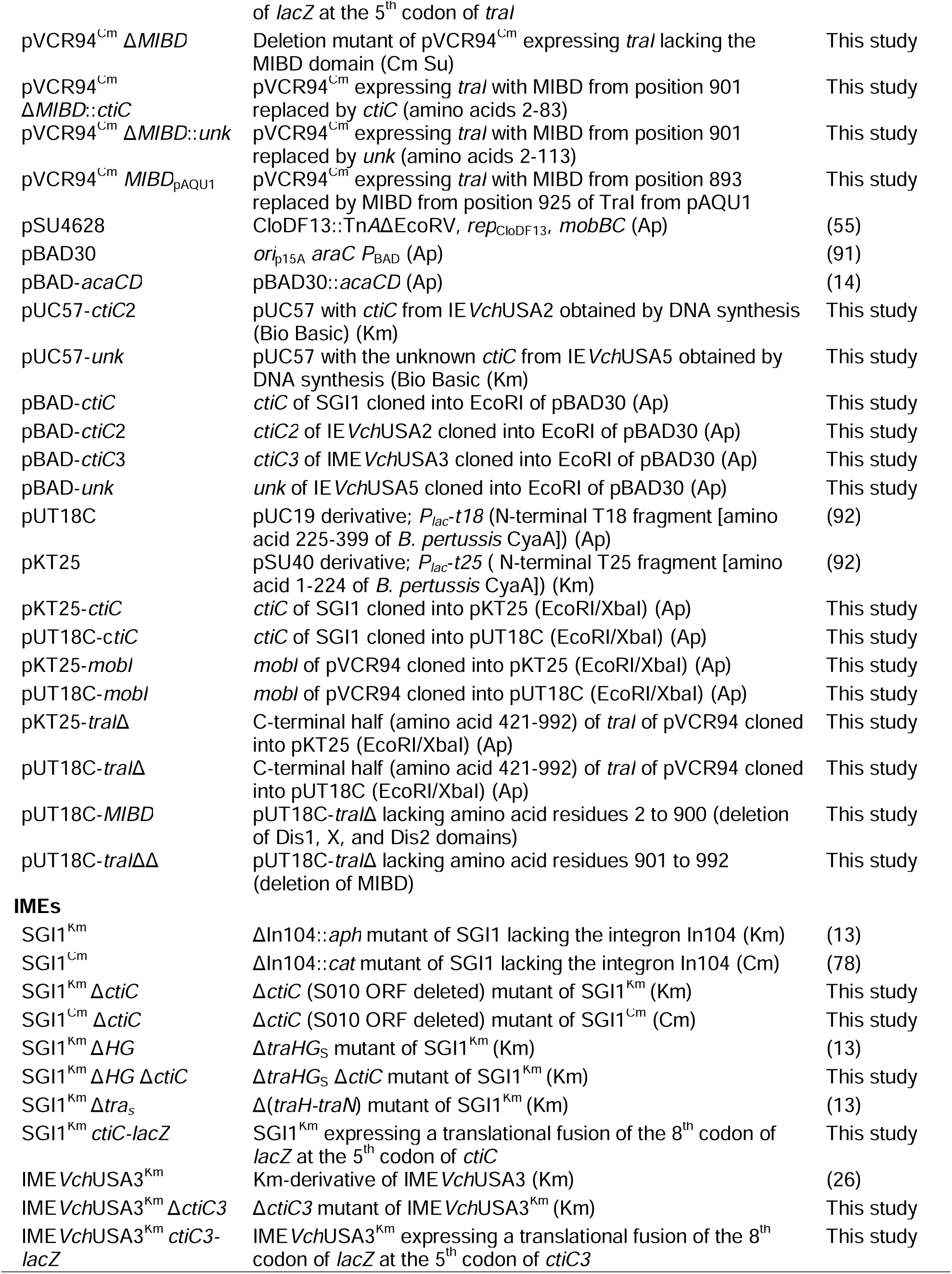
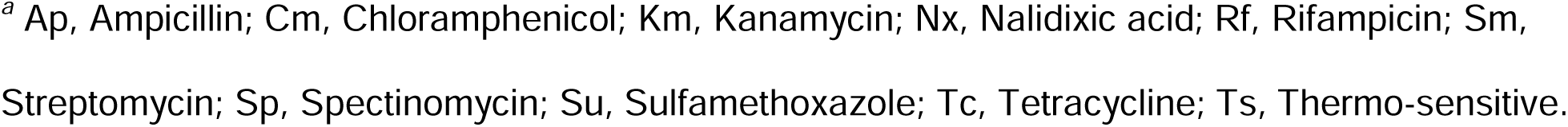
Strains, plasmids and IMEs used in this study.

### Conjugation assays

Except for the TraDIS experiment (see below), conjugation assays were performed by mixing equal volumes of donor and recipient cultures grown at 37°C with shaking for 4 hours. The cells were harvested by centrifugation, washed in 1 volume of LB broth and resuspended in 1/20 volume of LB broth. The mixtures were then deposited on LB agar plates and incubated at 37°C for 2 hours. After mating, the cells were recovered from the plates in 1 ml of LB broth, and serial dilutions were prepared in LB broth for immediate plating. Donors, recipients and transconjugants were selected on LB agar plates containing appropriate antibiotics. Frequencies of transfer were calculated by dividing the number of transconjugant CFUs by the number of donor CFUs. When required for induction of gene expression during complementation experiments, LB agar plates were supplemented with L-arabinose (0.02%, wt/vol).

### Molecular biology methods

Genomic DNA and plasmid extraction was prepared using QIAamp DNA mini kit and QIAprep Spin miniprep kit (Qiagen) or Monarch Plasmid Miniprep Kit (New England Biolabs), respectively, according to the manufacturer’s instructions. Restriction enzymes, Antarctic Phosphatase and T4 DNA Ligase used in this study were purchased from New England Biolabs. Several DNA polymerases were used: Q5 (New England Biolabs), Taq (New England Biolabs) and EasyTaq (Civic Bioscience). PCR products were purified using the QIAquick PCR Purification Kit (Qiagen), according to manufacturer’s instructions. *E. coli* was transformed by electroporation as described by Dower *et al.* (32) in a Bio-Rad GenePulser Xcell apparatus set at 25 µF, 200 Ω and 1.8kV using 1-mm electroporation cuvettes. Sanger sequencing reactions were performed by the Plateforme de Séquençage et de Génotypage du Centre de Recherche du CHUL (Québec, QC, Canada).

### Plasmid and strain construction

Plasmids and oligonucleotides used in this study are listed in Table 1 and Supplementary Table S1. Deletion and insertion mutants were constructed using the one-step chromosomal gene inactivation technique of Datsenko and Wanner (33). All deletions were designed to be non-polar. *traHG* and *tra_S_* deletion mutants of SGI1^Km^ were constructed previously (13). Details about constructs are provided in Supplementary Text S1.

### Transposon-directed insertion sequencing (TraDIS)

A conjugation-assisted random transposon mutagenesis experiment was performed on *E. coli* KH95 bearing pVCR94^Km^ Δ*acr2 trmE*::SGI1^Cm^ using the transposition system of *E. coli* MFD*pir+* carrying pFG036 and pFG051 as described previously (15). Details are provided in Supplementary Text S1.

### Preparation of TraDIS libraries and Illumina sequencing

For each library, a 1.5 ml frozen stock aliquot was thawed on ice for 15 min and used to prepare sequencing libraries as described previously (15). Mutant libraries were then pooled and sequenced by Illumina using the NextSeq 500/550 High Output Kit v2 at the RNomics platform of the Laboratoire de Génomique Fonctionnelle de l’Université de Sherbrooke (https:// rnomics.med.usherbrooke.ca) (Sherbrooke, QC, Canada). The transposon data analysis was carried out as described previously (15).

### Bacterial adenylate cyclase two-hybrid (BACTH) assays

Protein-protein interactions were examined using a bacterial two-hybrid assay based on the reconstitution of the adenylate cyclase CyaA of *Bordetella pertussis* from the fragment T18 and T25 expressed from the two BACTH vectors pUT18C and pKT25 (34). The plasmid constructs expressing the genes of interests were transformed into *E. coli* BTH101. CyaA activity was measured through the activation of the expression of the *lacZ* reporter gene.

### β-galactosidase assays

β-galactosidase assays were carried out on LB agar plates supplemented with 40 μg/mL 5-bromo-4-chloro-3-indolyl-β-D-galactopyranoside (X-gal) and 0.5 mM isopropyl β-D-1-thiogalactopyranoside (IPTG) for 36h at room temperature or in LB broth using *o*-nitrophenyl-β-D-galactopyranoside (ONPG) as the substrate as described previously (35). Overnight cultures were diluted 1:100, and further grown to mid-log phase (OD_600_ = 0.6). The mean values and standard error of the mean of β-galactosidase activities were calculated from 3 independent experiments.

### RNA extraction and cDNA synthesis

RNA extractions and cDNA synthesis were performed as previously described (21) with the following modifications. *E. coli* VB112 containing SGI1^Km^ with or without pBAD-*acaCD* was grown at 37°C for 16 h in LB broth containing the appropriate antibiotics. cDNA was synthesized from 0.5 ng of RNA and 1 pmol of gene-specific primer ctiC_RT (Integrated DNA Technologies), using the reverse transcriptase SuperScript IV (Invitrogen), according to manufacturer’s instructions. Control reactions without reverse transcriptase treatment (‘RNA’) were performed for each sample. PCR reactions aiming at amplifying *traG*-*ctiC* and *traH* were carried out using 50 pg of RNA or the corresponding amount of cDNA, or 0.5 ng of gDNA as the template, and primer pairs traG_i_F2/ctiC_i_R and traH_i_F/traH_i_R, respectively.

### Multiple sequence alignment for sequence comparison and phylogenetic analyses

Initial comparative analyses of CtiC were conducted with HHblits (https://toolkit.tuebingen.mpg.de/tools/hhblits) (36). The diversity of domain architectures of proteins related to CtiC was assessed with a PSI-Blast analysis of CtiC against the NCBI Protein Reference Sequences restricted to Gammaproteobacteria (5 iterations, taxid: 1236). The recovered sequences where clustered using mmseqs2 (https://toolkit.tuebingen.mpg.de/tools/mmseqs2) with a minimum sequence identity parameter set to 1 to remove redundant sequences (37). Multiple sequence alignments (MUSCLE), alignment curation (trimAl) and tree inference with model selection and bootstrap (PhyML +SMS) were conducted as a workflow on the https://ngphylogeny.fr/ server (38–43). Trees were displayed and annotated with iTOL (44).

### Protein structure predictions

Modeling of protein structures was done using Alphafold2 with default parameters (45). The generated PDB files were visualized and analyzed using the ChimeraX 1.10 software (46). Structure similarity searches were performed by the protein structure comparison tools PDBeFold v2.58 (http://www.ebi.ac.uk/msd-srv/ssm) (47) and Dali (http://ekhidna2.biocenter.helsinki.fi/dali/) (48) against the whole PDB archive. All against all structure comparisons were computed on the Dali server. Sequence similarities and secondary structure information from aligned sequences were rendered using ESPript 3.0 (https://espript.ibcp.fr) (49). Interprotein interactions in AlphaFold2 predicted structures were scored by the Python script ipsae.py (https://github.com/DunbrackLab/IPSAE) (50).

### Statistical analyses and figures preparation

Numerical data presented in graphs are available in the Supplementary Dataset. All analyses were performed using R Statistical Software (v4.5.2) and RStudio (v2026.01.0.392) (51). We assumed a normal distribution of the frequencies of transconjugant formation and tested the homogeneity of variance of the log_10_-transformed frequencies using Bartlett’s tests. Dunnett and Tukey-Kramer tests and compact letter display were performed via the multcomp R package (v1.4-29) (52). Graphics were rendered via the ggplot2 (v4.0.1) (53) and pheatmap (v1.0.13) (54) R packages. All figures were prepared using Inkscape 1.4 (https://inkscape.org/).

## Results

### Identification of SGI1 genes that inhibit IncC plasmid transfer

Since SGI1 strongly inhibits A/C plasmid transfer, we hypothesized that disruption of SGI1 inhibitory genes should restore wild-type transfer of the IncC plasmid pVCR94 and increase the population of transconjugants harbouring both SGI1 and pVCR94. Hence, we performed a transposon-directed insertion sequencing (TraDIS) assay designed to enrich SGI1 mutants that allowed the cotransfer and establishment of pVCR94 in recipients (Fig. 1A). We constructed a library of high-density random mini-Tn*5* insertions in *E. coli* KH95 bearing pVCR94^Km^ Δ*acr2* and SGI1^Cm^. Removal of *acr2*, the primary repressor of A/C plasmid transfer, increases conjugative transfer, preventing an insertion bias in this negative regulator gene. The resulting mutants formed the input library enriched in mini-Tn*5* insertions that enable plasmid replication and coexistence with SGI1. We used the input library in a mating assay to transfer the transmissible set of mini-Tn*5*-bearing plasmids and SGI1^Cm^ to *E. coli* GG56 (Nx^r^). We then selected transconjugants that had acquired both pVCR94^Km^ and SGI1^Cm^ to make up the output library. High-throughput Illumina sequencing was used to map the mini-Tn*5* insertion sites in both the input and output libraries. An excess of insertions in the output compared to the input was expected to reveal IncC plasmid or SGI1 genes that inhibit the cotransfer.

**Figure 1.**
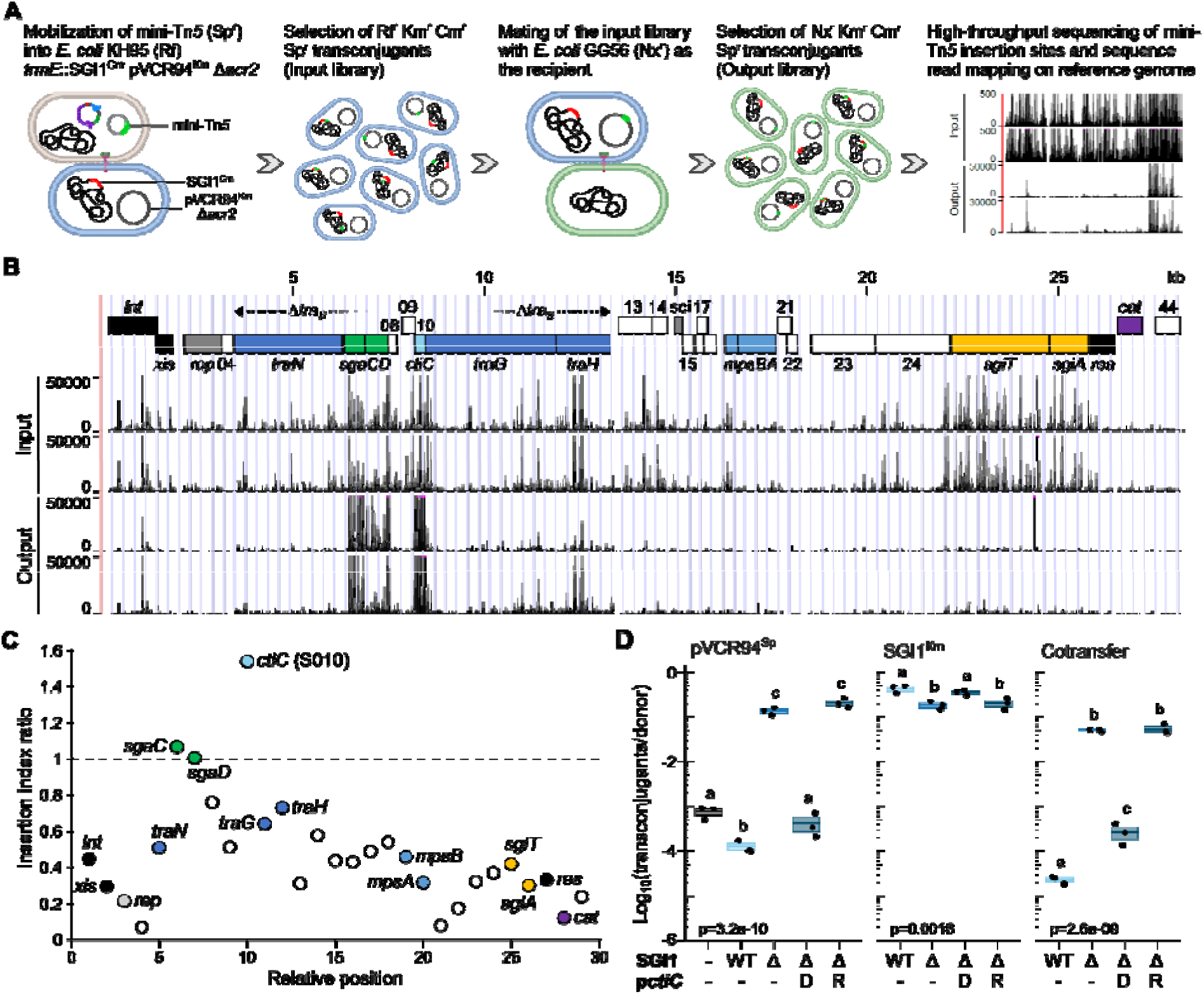
Identification of SGI1 genes inhibiting IncC plasmid transfer. (**A**) TraDIS strategy for identifying genes that inhibit the cotransfer of SGI1 and its helper IncC plasmids. Created with BioRender.com. (**B**) Read mapping of mini-Tn*5* insertions in SGI1 in *E. coli* KH95 (Rf^r^) bearing pVCR94^Km^ Δ*acr2 trmE*::SGI1^Cm^ (Input) and in *E. coli* GG56 (Nx^r^) transconjugants selected for cotransfer of pVCR94^Km^ and SGI1^Cm^ (Output). Genes on the map of SGI1^Cm^ (top track) are colour-coded as follows: black, recombination and transposition; dark grey, replication/partition; dark blue, mating pair formation; light blue, DNA processing; green, transcriptional activator; red, transcriptional repressor; yellow, toxin-antitoxin system; purple, antibiotic resistance; white, unknown function. In the input and output, each track corresponds to a biological replicate. (**C**) Dot plot of mini-Tn*5* insertion index for each gene (S0xx) of SGI1^Cm^. Insertion indices were calculated as the ratios of insertion counts between input and output. (**D**) Effect of *ctiC* (S010) on the conjugative transfer of pVCR94^Sp^. For all mating assays, donors are *E. coli* GG56 (Nx^r^) containing pVCR94^Sp^, and recipients are *E. coli* CAG18439 (Tc^r^). When indicated, the donor strain harboured SGI1^Km^ (WT) or its Δ*ctiC* mutant (Δ) with or without pBAD-*ctiC* (p*ctiC*). Expression of *ctiC* from pBAD30 in donor (D) or recipient (R) cells was induced using 0.02% arabinose. Crossbars show the mean and standard errors of the mean of at least three independent experiments. A one-way ANOVA with a Tukey-Kramer post hoc test was used to compare the means of the log_10_-transformed values. The compact letter display shows statistical significance for pairwise comparisons, where means grouped under identical letters are not statistically different. The complete set of *p*-values is available in Supplemental Dataset S1.

Before all else, we could not detect any statistically significant mini-Tn*5* insertion enrichment in the output for pVCR94^Km^, suggesting that no gene of the helper plasmid harms the mobilization of SGI1 in our conditions. In contrast, the 2.1-kb region of SGI1 bordered by *traN* and *traG* contained the bulk of mini-Tn*5* insertions (Fig. 1B). More specifically, a small open reading frame of unknown function, S010, was the most frequently disrupted gene (Fig. 1B and 1C). For this reason, S010 was renamed *ctiC* for <u>c</u>onjugative transfer inhibitor of Inc<u>C</u>. To confirm the involvement of *ctiC* in transfer inhibition, we performed a 2-h *E. coli* mating assay using donors bearing pVCR94^Sp^ and SGI1^Km^ or its Δ*ctiC* mutant. Whereas the presence of SGI1^Km^ reduced the transfer of pVCR94^Sp^ by 6-fold, deleting *ctiC* increased pVCR94^Sp^ transfer by ∼1,096-fold and the cotransfer by ∼2,186-fold (WT vs Δ, Fig. 1D). This deletion reduced the mobilization of SGI1^Km^ by 2.3-fold. Surprisingly, pVCR94^Sp^ transfer was enhanced (180-fold) in the presence of SGI1 Δ*ctiC*, reaching a level comparable to the IME (- vs Δ, Fig. 1D). This observation indicates that SGI1 improves the transfer of IncC plasmids if *ctiC* is missing.

The Δ*ctiC* mutation was complementable only upon ectopic expression of *ctiC* in the donor, not the recipient. The complementation inhibited IncC plasmid transfer and the cotransfer, albeit not as strongly as wild-type SGI1 (Fig. 1D). It also restored the transfer of SGI1^Km^ Δ*ctiC* to its wild-type level. Altogether, these results indicate that the product of *ctiC* acts by interfering with a conjugative process occurring in donor cells, not during the establishment of the plasmid in the recipient cell.

### CtiC specifically affects the transfer of A/C plasmids

Given the relatedness of IncC and IncA plasmids, we tested whether *ctiC* also affects the transfer of the IncA plasmid pRA1. The presence of SGI1 resulted in an 80-fold reduction of pRA1 transfer in a *ctiC*-dependent manner (Fig. 2A). The deletion of *ctiC* reduced SGI1 mobilization by 4-fold, while restoring the wild-type transfer level of pRA1. However, SGI1^Km^ Δ*ctiC* did not further improve the transfer of pRA1, unlike what we observed for pVCR94.

**Figure 2.**
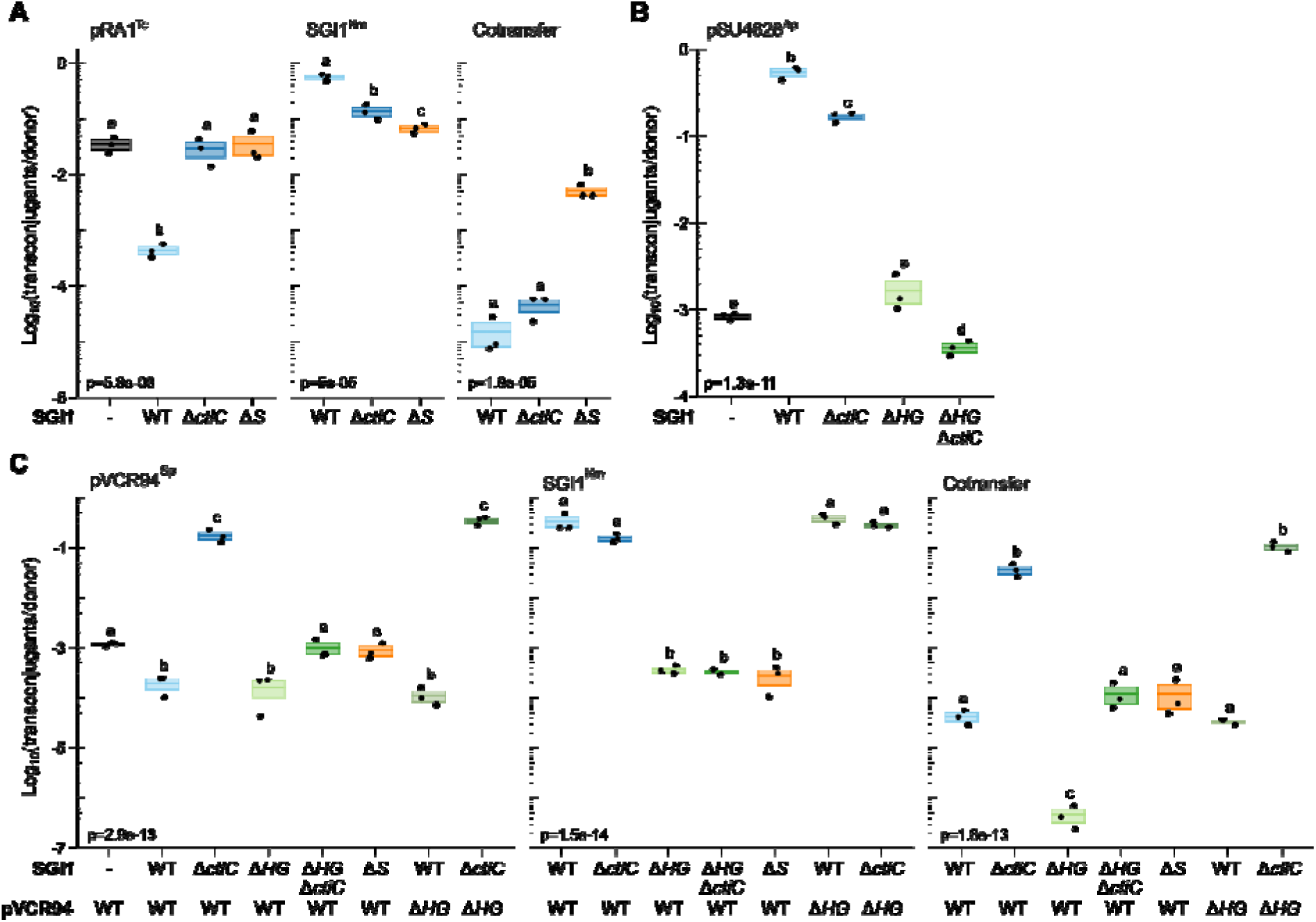
Effect of SGI1 and *ctiC* on conjugative transfer. (**A**) *ctiC* inhibits the transfer of the IncA plasmid pRA1. Donors are *E. coli* GG56 (Nx^r^) containing pRA1 (Tc) and SGI1^Km^ (WT) or its Δ*ctiC* mutant (Δ). Recipients are *E. coli* VB112 (Rf^r^). Transconjugants were selected as Rf^r^ Tc^r^ (pRA1), Rf^r^ Km^r^ (SGI1), or Rf^r^ Tc^r^ Km^r^ (Cotransfer) colonies. (**B**) Effect of SGI1 on the mobilization of pSU4628. Donors are *E. coli* GG56 (Nx^r^) containing pVCR94^Sp^ and pSU4628^Ap^, with or without (-) SGI1^Km^ (WT) or its designated mutants. (**C**) Effect of *traHG* of SGI1 on IncC plasmid transfer in the presence or absence of *ctiC*. Donors are *E. coli* GG56 (Nx^r^) containing pVCR94^Sp^ or its Δ*traHG* mutant and with or without (-) SGI1^Km^ (WT) or its designated mutants. In (A) and (B), the recipient is *E. coli* CAG18439 (Tc^r^). Transconjugants were selected as the Tc^r^ Ap^r^ (pSU4628), Tc^r^ Sp^r^ (pVCR94), Tc^r^ Km^r^ (SGI1), or Tc^r^ Sp^r^ Km^r^ (Cotransfer) colonies. Statistical analysis is as described in Fig. 1D. The complete set of *p*-values is available in Supplemental Dataset S1.

To determine whether *ctiC* affects the T4SS or the processing of A/C plasmid DNA, we tested its impact on the mobilization of pSU4628, an ampicillin-resistant derivative of pCloDF13, which is mobilizable by several types of conjugative plasmids (IncF, I, N, P, W, and C) (13, 55). pCloDF13 encodes two proteins, MobB and MobC, which are required for processing its *oriT* to allow mobilization (56). MobC is the relaxase (MOB_C_ family), whereas MobB, necessary for efficient *oriT* cleavage, acts as a type IV coupling protein. We used pVCR94^Sp^ with SGI1^Km^ or its Δ*ctiC* mutant to mobilize pSU4628^Ap^. pSU4628^Ap^ mobilization increased by more than 2 logs in the presence of SGI1 and was even strengthened by *ctiC* (650-fold for wild-type SGI1 vs 204-fold for SGI1 Δ*ctiC*) (Fig. 2B). Hence, CtiC does not impair the DNA processing functions or the type IV coupling protein of pCloDF13 or the T4SS, suggesting it affects A/C plasmids specifically. It also confirms that SGI1 enhances the conjugative properties of donor cells in some way. We hypothesize that this phenotype could result from the TraHG subunit substitution operated by SGI1 in the T4SS encoded by A/C plasmids (13).

### CtiC counteracts the effect of TraHG substitution, which increases IncC plasmid transfer

TraH is involved in mating pair formation/stabilization, whereas TraG is an inner membrane protein involved in mating pair stabilization and entry exclusion (20). To test whether TraH and TraG substitution enhanced conjugative transfer, we used SGI1^Km^ mutants lacking *traHG*, *ctiC*, both, or the region encompassing *traN* to *traH* (mutant Δ*tra_S_*, Fig 1B). The deletion of *traHG* in SGI1^Km^ seemingly had no impact on pVCR94^Sp^ transfer, which remained low due to *ctiC*; however, it reduced SGI1^Km^ transfer by 3 logs and nearly abolished cotransfer (WT vs Δ*HG*) (Fig. 2C). SGI1^Km^ Δ*traHG* also reduced the mobilization of pSU4628^Ap^ to the level of cells lacking SGI1 (Fig. 2B).

In the presence of SGI1^Km^, pVCR94^Sp^ and its Δ*traHG* mutant transferred at the same frequency, suggesting that *traHG* of SGI1 compensates the Δ*traHG* mutation; yet, even in the absence of the IncC *traHG* genes, SGI1^Km^ transferred at its highest frequency (Fig 2C), confirming the involvement of *traHG* of SGI1 in reaching high levels of transconjugant formation. The suppression of both *traHG* in pVCR94^Sp^ and *ctiC* in SGI1 resulted in very high transfer frequencies for the helper plasmid, SGI1 and cotransfer of both, indicating that SGI1 completely overrides the need for *traHG* of pVCR94 to enhance conjugative transfer.

In contrast, SGI1^Km^ Δ*traHG* Δ*ctiC* failed to enhance the transmissibility of pVCR94^Sp^. However, it restored pVCR94^Sp^ transfer to wild-type level (Fig. 2C). SGI1^Km^ Δ*traHG* and SGI1^Km^ Δ*traHG* Δ*ctiC* transferred at comparable low levels. Moreover, SGI1^Km^ Δ*traHG* Δ*ctiC* led to the lowest mobilization frequency of pSU4628*Ap*, presumably due to the three mobile elements competing for access to the T4SS (Fig. 2B and 2C). Finally, using pVCR94^Sp^ as the helper plasmid, all transfer frequencies using SGI1^Km^ Δ*traHG* Δ*ctiC* were indistinguishable from those obtained using SGI1^Km^ Δ*tra_S_*, which lacks *traN*, *sgaDC*, *ctiC*, two open reading frames of unknown function, and *traHG* (Fig. 2C). These results indicate that TraH and TraG of SGI1 are necessary to enhance the conjugative properties of the T4SS encoded by IncC plasmids. Since SGI1^Km^ Δ*ctiC* failed to boost pRA1^Tc^ transfer, we hypothesize that *traHG* of SGI1 does not improve upon the functionality of the *traHG* homologues carried by the IncA plasmid (Fig. 2A). This hypothesis is supported by the SGI1^Km^ Δ*tra_S_* mutant, which, despite lacking both *ctiC* and *traHG*, was mobilized at a frequency 250 times higher by pRA1^Tc^ than by pVCR94^Sp^ (Fig. 2A and 2C).

Altogether, our observations suggest that *ctiC* specifically targets A/C plasmids and prevents them from benefiting from an enhancement of conjugative transfer mediated at least in part by the substitution of TraH and TraG subunits in the A/C-encoded T4SS. Next, we focused on the mechanistic aspect of the transfer inhibition mediated by CtiC.

### CtiC resembles the C-terminal domain of MOB_H1_ relaxases

The translation product of *ctiC* is an 84-amino acid residue, basic protein (pI=10.08) with a predicted winged helix fold resembling the archaeal DNA polymerase holoenzyme subunit PBP2 (PolB1 binding protein 2) (Supplementary Fig. S1A) (57). The topology of the winged helix fold consists of a bundle of three α-helices and two wings of a 3- or 4-strand β-sheet (wing) (58, 59). The surface of CtiC formed by the first half of the helix α1 and β-sheet has a net positive charge, whereas the opposite surface is negatively charged (Supplementary Fig. S1C). Comparative analysis of CtiC using Blast and HHblits revealed two types of proteins.

First, *ctiC* is conserved in most SGI1-related IMEs integrated at the 3’ end of *trmE*, including GI-15 of the seventh pandemic *V. cholerae* O1 El Tor, PGI2 of *P. mirabilis*, AGI1 of *A. baumannii*, and KGI1 of *K. pneumoniae* (Fig. 3A and Supplementary Fig. S2). IMEs integrated at the 5’ end of *dusA* (e.g., IME*Vch*USA3) or the 3’ end of *yicC* (e.g., IE*Sen*USA1) encode highly divergent CtiC homologues (24 to 33% identity with CtiC of SGI1) (Fig. 3A and 3B). As in SGI1, their coding sequence abuts *traG* except for type 3 *dusA*-associated IMEs (e.g., *unk* of IE*Vch*USA5), where it is separated from *traG* by a small ORF of unknown function (Supplementary Fig. S2). The evolutionary relationship between the CtiC homologues is congruent with that of their cognate IMEs (26). Comparison of the predicted 3D structures revealed a similar fold for CtiC, CtiC2 and CtiC3, except for an additional N-terminal α helix in the latter two (Fig. 3A). In contrast, Unk of type 3 IMEs folds into a different conformation (Fig. 3A and 3C).

**Figure 3.**
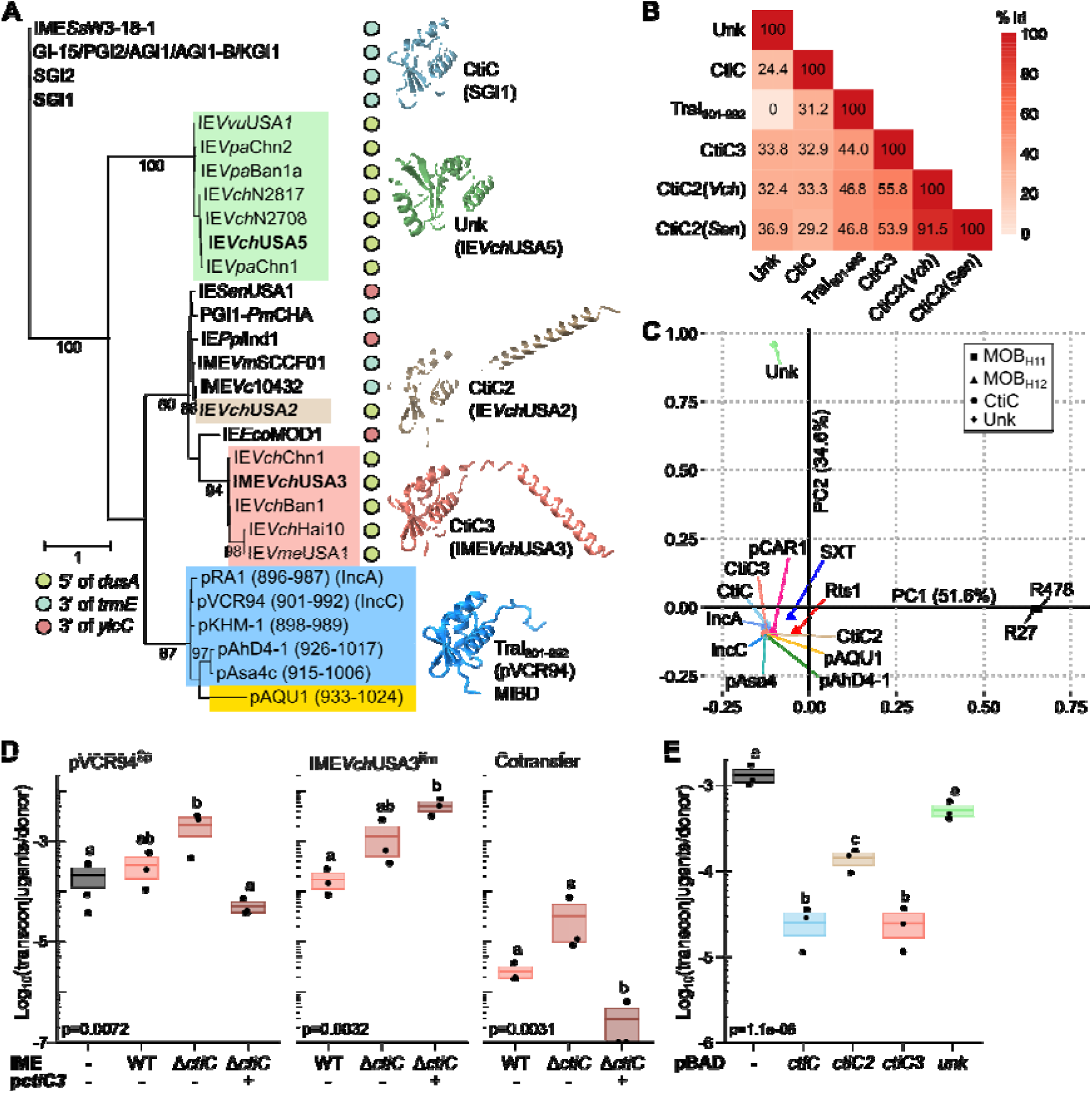
CtiC-related proteins or domains. (**A**) Maximum likelihood phylogenetic analysis of CtiC homologues. Bootstrap supports are indicated as percentages at the branching points only when >80%. Branch lengths represent the number of substitutions per site over 83 sites using the HIVb substitution model +G. Colored disks indicate integration sites. Lineages with pink, tan, and green backgrounds indicate *dusA*-integrated IMEs of type 1, 2, and 3, respectively, as defined previously (26). The lineage with the blue and gold backgrounds contains relaxase CTDs with the predicted structure of TraI_C_92_ used as the reference. The predicted structures of CtiC homologues encoded by SGI1, IE*Vch*USA2, IME*Vch*USA3, and IE*Vch*USA5 are shown. (**B**) Pairwise comparison heatmap of blastp identity percentages of CtiC homologues and TraI_901-992_. Protein accession numbers are as follows: CtiC (AAK38390.1), CtiC3 (WP_095463402.1), CtiC2(*Vch*) (OFI96794.1), CtiC2(*Sen*) (WP_069140155.1), Unk (WP_095467696.1), TraI_901-992_ (WP_011872892.1). (**C)** Structural similarity correspondence analysis of representative CtiC homologues and CTD of MOB_H1_ relaxases computed by the Dali server. (**D**) and (**E**), 2-h mating assays between *E. coli* GG56 (Nx^r^) containing pVCR94^Sp^ (donor) and *E. coli* CAG18439 (Tc^r^) (recipient). Transconjugants were selected as Tc^r^ Sp^r^ (pVCR94), Tc^r^ Km^r^ (IME*Vch*USA3), or Tc^r^ Sp^r^ Km^r^ (Cotransfer) colonies. In D, donors harboured IME*Vch*USA3^Km^ (WT) or its Δ*ctiC3* mutant with (+) or without (-) pBAD30 expressing *ctiC3* (p*ctiC3*). In E, the donor expressed the designated *ctiC* from pBAD30. Crossbars show the mean and standard errors of the mean of three independent experiments. Statistical analyses are as described in Fig. 1D. The complete set of *p*-values is available in Supplemental Dataset S1.

Second, CtiC is distantly related to the last 92 amino acid residues of the C-terminal domain (CTD) of the relaxase TraI of A/C plasmids (31.2% identity and 51% similarity) (Fig. 3B). A PSI-Blast analysis revealed that the CtiC signature motif exists only in two protein architectures, either as an autonomous entity or associated with a TraI_2 relaxase domain (Pfam PF07514) (Supplementary Fig. S3 and Supplementary Table S2). CTDs of MOB_H12_ relaxases encoded by distant relatives of A/C plasmids (pAQU1, pAsa4 and pAhD4-1), IncT (Rts1) and IncP-7 (pCAR1) plasmids, and SXT/R391 elements adopt a similar conformation (Supplementary Fig. S4A and S4C). CTDs of MOB_H11_ relaxases from IncHI1 (R27) and IncHI2 (R478) plasmids are more dissimilar (Fig. S4B and S4C). CTDs of relaxases encoded by A/C and related plasmids form a clade that is distinct from CtiC homologues encoded by SGI1-related IMEs, regardless of their integration site (Fig. 3A). However, the origin of CtiC homologues cannot be clearly ascribed to a single common ancestor (Supplementary Fig. S3). Correspondence analysis of all predicted structures confirmed that, except Unk from type 3 IMEs and MOB_H11_ relaxase CTDs, CtiC homologues and MOB_H12_ relaxase CTDs have almost identical conformations (Fig. 3C and Supplementary Fig. S4C).

### Distantly related CtiC homologues inhibit IncC plasmid transfer

Given the divergence of SGI1-related IMEs inserted at *dusA*, we wondered whether the CtiC homologues they encode could inhibit IncC plasmid transfer. First, we tested the effect of IME*Vch*USA3^Km^ or its Δ*ctiC3* mutant on pVCR94^Sp^ transfer. Deleting *ctiC3* slightly increased (>6-fold) plasmid transfer but had no significant impact on IME mobilization (Fig. 3D). Notably, IME*Vch*USA3 lacks *traH* (Supplementary Fig. S2), which may explain the limited gain in conjugative transfer compared to SGI1. Overexpressing *ctiC3* in donors reduced plasmid transfer by >6-fold and increased IME mobilization by ∼30-fold.

Next, we compared the inhibitory activity of *ctiC* of SGI1, IE*Vch*USA2, IME*Vch*USA3, and the distant homologue *unk* of IE*Vch*USA5 in donors bearing pVCR94^Sp^. *ctiC* genes reduced pVCR94^Sp^ transfer of pVCR94^Sp^ by 38- to 126-fold (Fig. 3E). In contrast, *unk* had no impact. Hence, all but the most distant CtiC homologues inhibit IncC plasmid transfer.

### CtiC reveals a MobI-binding domain (MIBD) conserved in MOB_H12_ relaxases

Based on the predicted structural similarity between CtiC and the CTD of TraI, we hypothesized that TraI’s CTD binds to the mobilization factor MobI and that CtiC interacts with MobI, precluding the recruitment of TraI to *oriT*. Thus, we examined possible interactions between CtiC, MobI, and TraI using a bacterial adenylate cyclase-based two-hybrid (BACTH) assay. The T18 or T25 fragments of the split-adenylate cyclase were fused to the N-terminus of each protein. Because cloning of full-length *traI* proved unsuccessful due to possible toxicity, we used TraIΔHD, which lacks the N-terminal TraI_2 HD hydrolase domain, for all fusions. We detected the self-association of MobI, suggesting it works as a dimer or multimer, but failed to observe CtiC or TraIΔHD self-association (Fig. 4). Likewise, CtiC and TraIΔHD did not interact. In contrast, we detected a high β-galactosidase activity for MobI/CtiC and a low activity for MobI/TraIΔHD, suggesting that MobI directly interacts with CtiC and TraI. These results support TraI’s CTD as a MobI-binding domain (hereafter named MIBD) and suggest that CtiC competes with TraI’s MIBD to bind to MobI.

**Figure 4.**
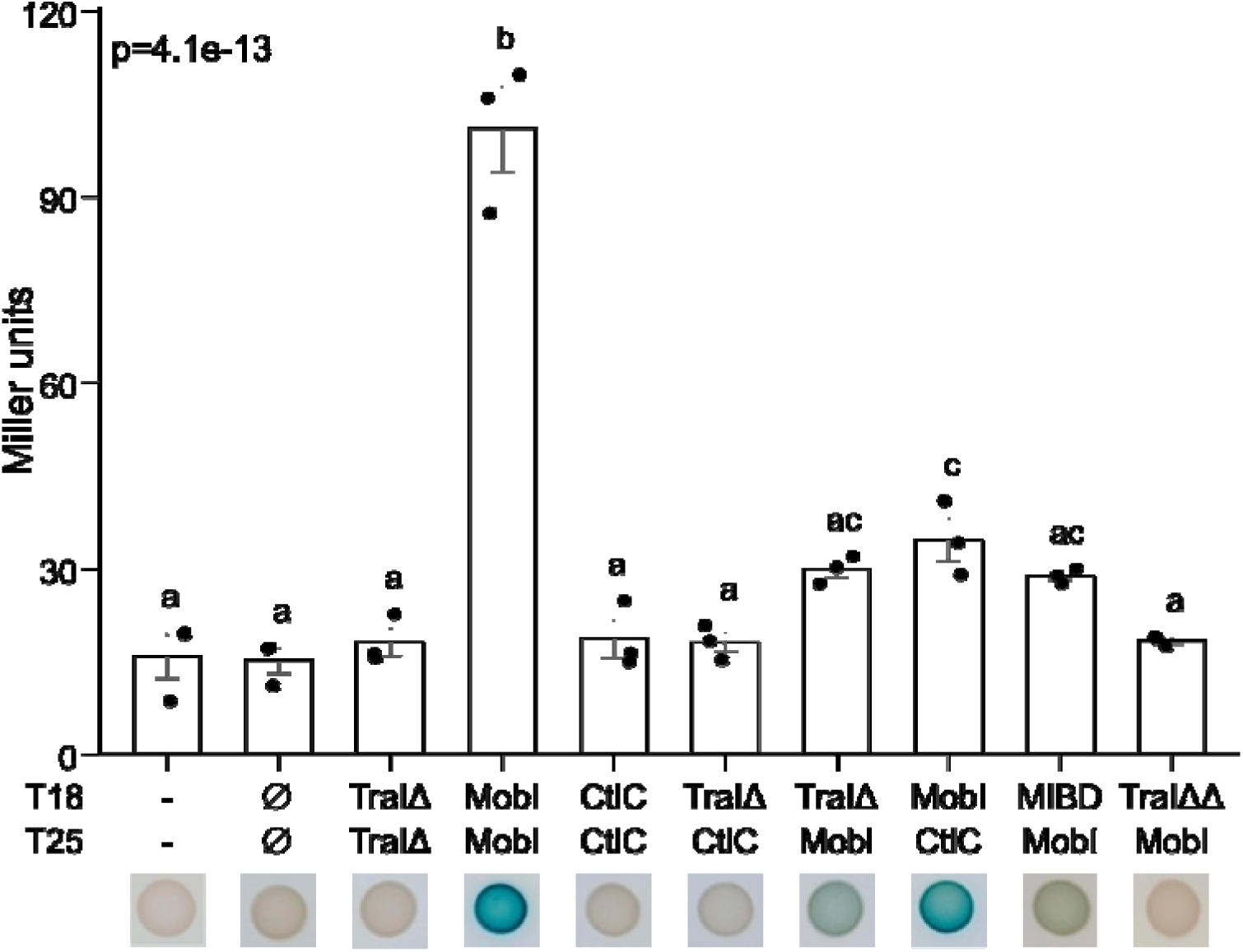
*In vivo* analysis of protein interactions using BACTH assays. MobI, TraI, MIBD and CtiC interactions were quantified in *E. coli* BTH101 carrying plasmids encoding T18 and T25 C-terminal fusions. “-” and “Ø” represent the negative controls BTH101 without plasmid and BTH101 carrying plasmids expressing unfused T18 and T25, respectively. TraIΔ lacks the TraI_2 HD hydrolase domain, whereas TraIΔΔ also lacks MIBD. Below the bars are shown the colonies grown on LB agar with IPTG. The bars represent the mean β-galactosidase activities and standard error of the mean of three independent experiments. A one-way ANOVA with a Tukey-Kramer post hoc test was used to compare the mean values. The complete set of *p*-values is available in Supplemental Dataset S1.

Next, we used BACTH assays to test whether MIBD is necessary and sufficient to promote the interaction with MobI using TraIΔHD mutants lacking MIBD and MIBD alone. We observed no β-galactosidase activity for the ΔMIBD mutant and low activity for MIBD alone, confirming that MIBD is required and sufficient for binding to MobI (Fig. 4).

### MIBD confers selectivity for MobI

Given the weak interactions between MobI and TraI, or its MIBD, observed in the BACTH assays, we made an MIBD substitution in *traI* to test the selectivity for MobI (Fig. 5A). First, we constructed a mutant of pVCR94^Cm^ expressing a chimeric *traI* gene where *ctiC* or *unk* replaced the coding sequence of MIBD. While the mutation *traI*ΔMIBD::*aph* and the substitution *traI*ΔMIBD::*unk* abolished pVCR94^Cm^ transfer, the substitution *traI*ΔMIBD::*ctiC* slightly restored it (Fig. 5D). In this case, transfer was detectable in a 2-h mating assay at a level ∼2,500 times lower than for wild-type *traI*. This result confirms the crucial role of MIBD in initiating conjugative transfer. It also indicates that CtiC can, albeit barely, substitute for MIBD, supporting that both MIBD and CtiC bind to the same target, MobI.

**Figure 5.**
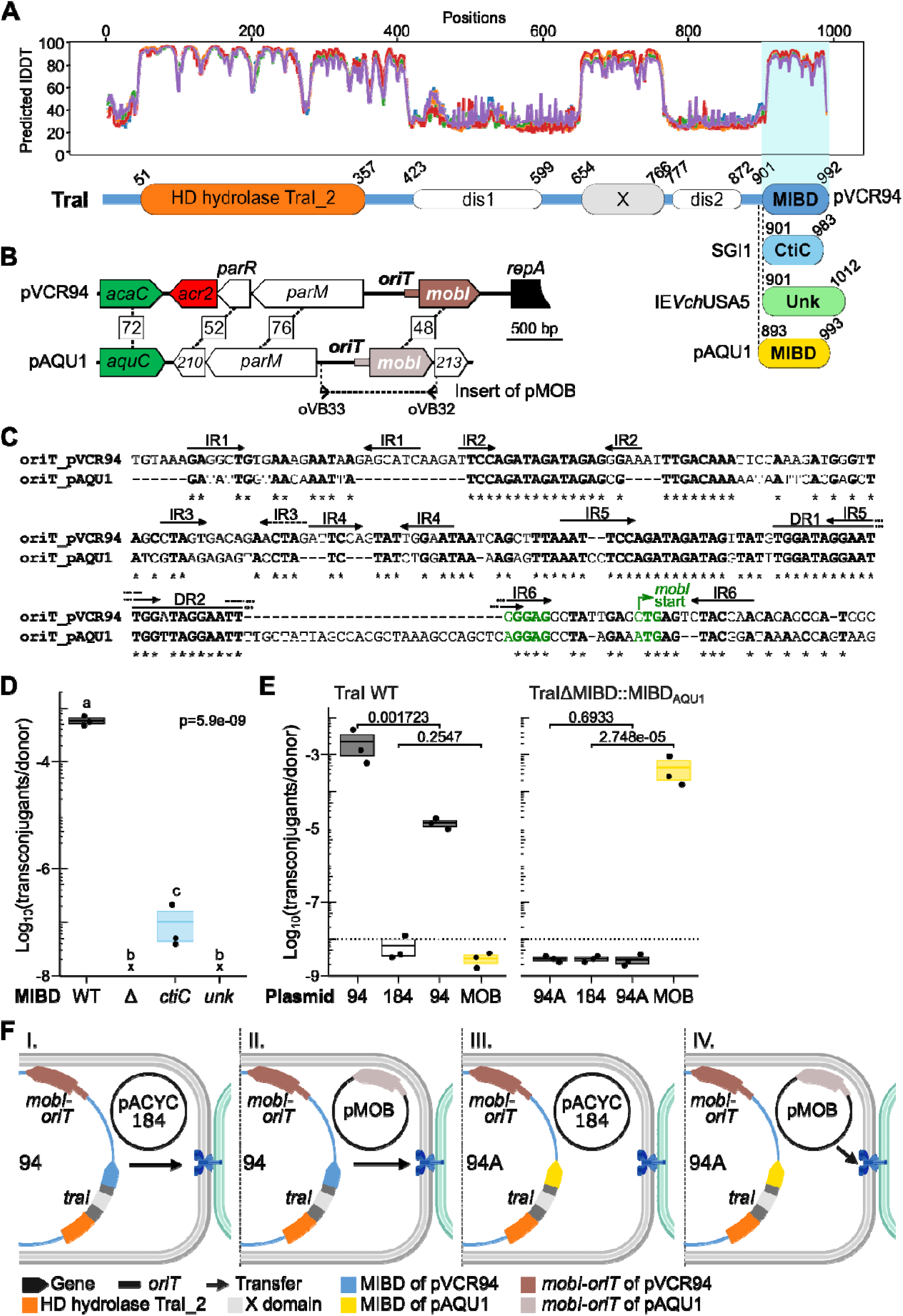
MIBD of TraI is necessary and sufficient for MobI binding. (**A**) Predicted local difference distance test (plDDT) plot of Alphafold2 structure predictions for pVCR94’s TraI. Higher plDDT implies higher accuracy of the prediction. Matching domains and consensus disorder predictions (Pfam and MobiDB-lite) made by InterProScan are shown below the plDDT plot. X and MIBD were added manually based on Alphafold predictions. Substitutions of the MIBD_VCR94_ domain by CtiC, Unk, or the MIDB domain of TraI of pAQU1 are indicated below at the indicated amino acid residue positions. (**B**) Comparison of the *oriT*-*mobI* region of pVCR94 and pAQU1. Boxed numbers on lines connecting two genes correspond to protein sequence identities. The region cloned into the non-mobilizable plasmid pACYC184Δ*cat2* to yield pMOB is shown by a dashed line, delimited by the primers used for PCR amplification. (**C**) MUSCLE alignment of the *oriT* loci of pVCR94 and pAQU1. Identical nucleotides are shown in bold and indicated by an asterisk. The inverted (IR) and direct (DR) repeats correspond to those identified by Hegyi *et al*. (19) in the *oriT* of IncC plasmids. (**D**) Substituting CtiC for MIBD partially complements the absence of MIDB. The recipient is *E. coli* CAG18439 (Tc^r^). Donors are *E. coli* GG56 (Nx^r^) containing pVCR94^Cm^ (WT) or its ΔMIBD (Δ), ΔMIBD-*ctiC* (*ctiC*), or ΔMIBD-*unk* (*unk*) *traI* mutants. Transconjugants were selected as the Tc^r^ Sp^r^ colonies. “x” indicates the frequency of transconjugant formation was below the detection limit (<10^-8^). Statistical analyses are as described in Fig. 1D. The complete set of *p*-values is available in Supplemental Dataset S1. (**E**) Substituting MIBD_AQU1_ for MIBD_VCR94_ changes the selectivity of TraI for the *oriT*-*mobI* module. The recipient is *E. coli* VB112 (Rf^r^). Donors are *E. coli* GG56 (Nx^r^) containing pACYC184Δ*cat2* (184) or pMOB (MOB) and pVCR94^Cm^ (94) or its *traI*ΔMIBD::MIBD_AQU1_ mutant (94A). Transconjugants were selected as the Rf^r^ Cm^r^ (pVCR94) or Rf^r^ Tc^r^ (pACYC184Δ*cat2* or pMOB) colonies. The *p*-values of unpaired Student’s t-tests are shown for each of the compared pairs. The dashed lines represent the detection limit (<10^-8^). (**F**) Schematic overview of the outcomes of panel 5E. The *traI* gene is represented in dark gray, with its specific domains color-coded according to the colors keys indicated in the legend. Successful transfer is indicated by a back arrow. I. and II., transfer of pVCR94^Cm^ wildtype only, supported by itself; III., no transfer; IV., transfer of pMOB only, supported by pVCR94^Cm^ *traI*ΔMIBD::MIBD_AQU1_.

Next, we replaced the MIBD coding sequence in *traI* of pVCR94^Cm^ with the corresponding sequence in *traI* of the conjugative plasmid pAQU1, a distant, untyped, relative of A/C plasmids (Fig. 5A). We also introduced into the donor strain pMOB, a derivative of the non-mobilizable plasmid pACYC184, within which we cloned the predicted *oriT*-*mobI* region of pAQU1 (Fig. 5 B). Although this region resembles that of pVCR94, its predicted *oriT* locus is highly divergent, while the predicted MobI shares only 48% identity (Fig. 5B and 5C). Mating assays revealed that pVCR94^Cm^ is unable to mobilize pMOB (Fig. 5E). Furthermore, the presence of *oriT*-*mobI* of pAQU1 even interfered with the transfer of pVCR94^Cm^ (160-fold reduction). In contrast, although pVCR94^Cm^ expressing the chimera *traI*ΔMIBD::MIBD_AQU1_ was unable to self-transfer, it mobilized pMOB at a high frequency, confirming that MIBD directly interact with MobI and confers the selectivity for the *oriT*-*mobI* locus independently of the HD_hydrolase TraI_2 domain of the relaxase.

Altogether, these observations demonstrate that MIBD and CtiC bind to the same partner, MobI. Although CtiC mimics the overall predicted structure of MIBD, it is a poor substitute when introduced into TraI and is suboptimal for initiating conjugative transfer.

Based on the results presented above, we hypothesized that CtiC inhibit A/C plasmid transfer by sequestering MobI, preventing its interaction with TraI. To do so, CtiC must be produced in amounts at least equivalent to MobI and act before TraI is expressed and binds to MobI. To test this hypothesis, we assessed the relative expression of *traI*, *mobI* and *ctiC*.

### Relative expression of TraI, MobI, and CtiC supports a competition model of fertility inhibition

*ctiC* abuts *traG*, suggesting it is the last gene of the *traHG* operon controlled by an AcaCD-responsive promoter (14, 60). To test this, we translationally fused a promoterless *lacZ* reporter gene to the fifth codon of *ctiC* in SGI1^Km^ and *ctiC3* in IME*Vch*USA3 (Fig. 6A). β-galactosidase assays showed that the transcriptional activator complex AcaCD, ectopically expressed from the arabinose-inducible promoter *P_BAD_*, increased the expression level of the *ctiC*’-*‘lacZ* fusions by ∼7- and ∼4-fold for SGI1 and IME*Vch*USA3, respectively (Fig. 6B). Furthermore, we detected a constitutive expression of *ctiC* at a level ∼4.5 times lower than that of *ctiC3* in non-inducing conditions. Next, we carried out a reverse transcription experiment using a primer located at the 3’ end of *ctiC* (ctiC_RT) and RNA obtained from *E. coli* VB112 cells containing SGI1^Km^ with or without pBAD-*acaCD* (Fig. 6A). PCR amplification using the generated cDNA as the template detected a weak transcript containing the junction between *traG* and *ctiC* (*traG*-*ctiC*) in the absence of *acaCD*, but did not detect a transcript containing *traH* (Fig. 6C). This observation confirms that *ctiC* is expressed constitutively from a weak promoter within *traG*. Furthermore, we observed a strong signal for both fragments upon expression of *acaCD*, which is consistent with *ctiC* being part of the *traHG* operon driven from the AcaCD-responsive promoter *P_traH_*.

**Figure 6.**
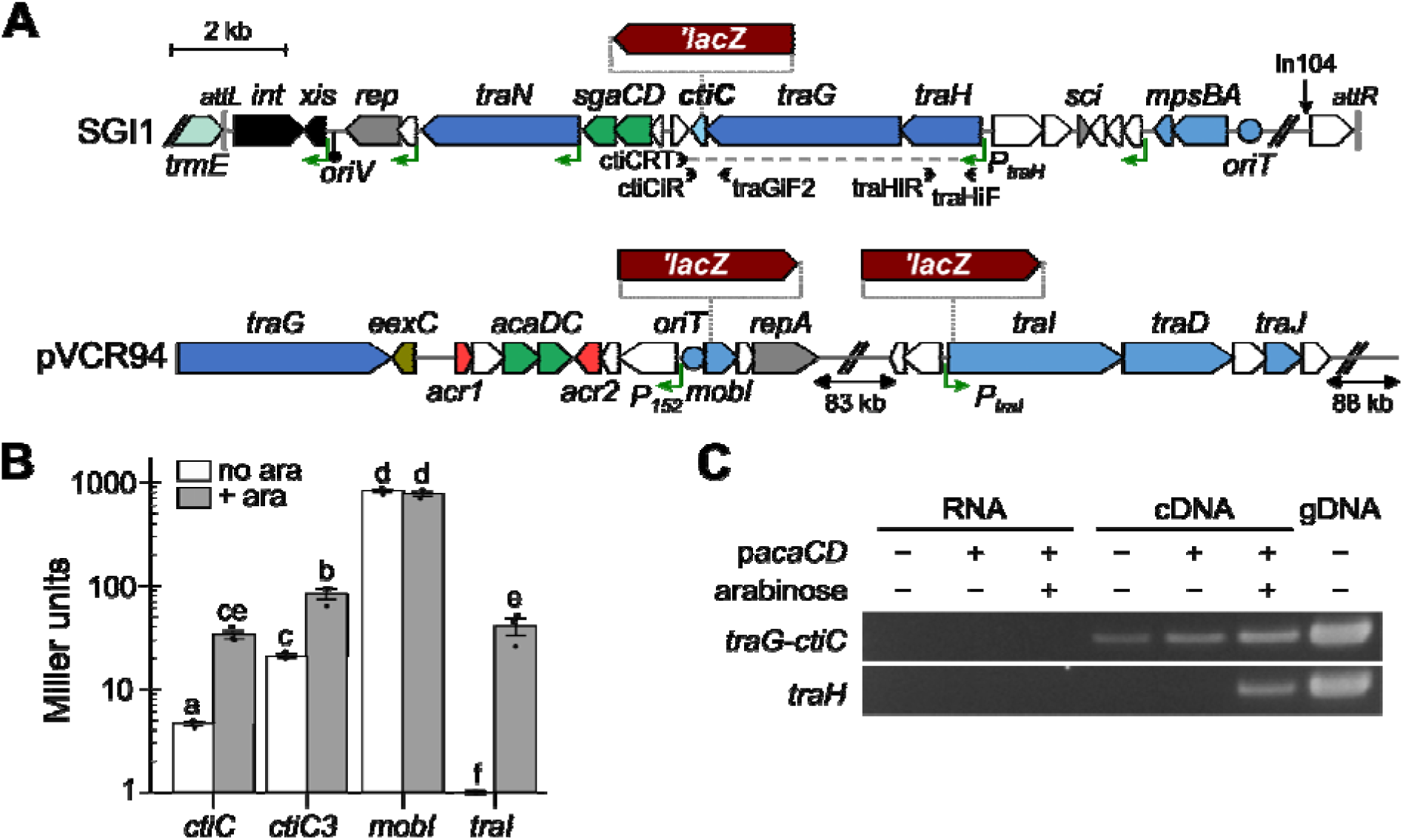
Comparative analysis of *ctiC*, *mobI*, and *traI* expression levels. (**A**) Schematic map of SGI1 and pVCR94 with the positions of the *lacZ* fusions. The *aad7* gene (Sp^r^), located downstream of *lacZ*, is not shown. Genes are colour-coded as indicated in Fig. 1. Green angled arrows show AcaCD-responsive promoters. Arrowheads show the position and name of primers used for the reverse transcription assays. (**B**) β-galactosidase activities expressed in Miller units of the *ctiC*’-, *ctiC3*’-, *mobI*’-, and *traI*’-’*lacZ* translational fusions in response to AcaCD in arabinose-induced (grey) versus non-induced (white) strain bearing pBAD-*acaDC*. The bars represent the mean and standard error of the mean of three independent experiments. Statistical analysis is as described in Fig. 1D. The complete set of *p*-values is available in Supplemental Dataset S1. (**C**) A 2% agarose gel of the PCR amplification of *traH_S_*and the junction between *traG_S_* and *ctiC* on the cDNA product. GG56 *trmE*::SGI1^Km^ genomic DNA (gDNA) and reverse transcription samples without reverse transcriptase (RNA) were used as positive and negative PCR controls, respectively.

We also translationally fused *lacZ* to the fifth codon of *traI* and *mobI* in pVCR94 (Fig. 6A) to quantify the expression of these two genes. *traI* expression was hardly detectable in the absence of *acaCD* and induced ∼300-fold in its presence (Fig. 6B). Expression levels of *traI*, *ctiC*, and *ctiC3* were comparable (40, 33, and 76 Miller units, respectively) in inducing conditions. In non-inducing conditions, *ctiC* and *ctiC3* surpassed *traI* expression by 34- and 160-fold, respectively. Finally, *mobI* expression was constitutive and topped the expression of all the other genes by 10- to 6,000-fold (Fig. 6B).

These findings support a model of fertility inhibition in which CtiC sequesters at least a fraction of the MobI molecules even before *traI* is expressed, thereby decreasing the likelihood of interactions between TraI and MobI. Hence, CtiC inhibits A/C plasmid transfer by competing with MIBD of TraI for recruiting MobI, thereby precluding TraI from binding to and processing the *oriT*.

## Discussion

Fertility inhibition is a process by which a plasmid impairs the conjugation of another co-resident plasmid in the donor cells (31). Various fertility inhibition systems target different steps of the conjugation process of coresident plasmids belonging to diverse incompatibility groups. For instance, FinO of IncFII plasmids inhibits the conjugation of the F plasmid by repressing the expression of its transfer genes via an increase in the antisense RNA FinP levels (61). Likewise, FinQ and FinU of IncI1 plasmids or FinW of IncFI plasmids downregulate the expression of F transfer genes via alternative pathways (62). FinV of IncX plasmids seems to impair the pilus assembly of IncF plasmids (63, 64). Other fertility inhibition systems affect the type IV coupling protein that connects the relaxosome to the T4SS. PifC of F targets TraG of the IncP plasmids, while FinC of pCloDF13 seems to target TraD of F (65, 66). The oncogenic suppressor Osa of the IncW plasmid pSa targets the Ti plasmid of *Agrobacterium tumefaciens* by degrading T-DNA attached to the VirD2 relaxase before it is translocated through the T4SS via the coupling protein VirD4 (67).

Here, we found that the IME SGI1 and kin (e.g., IME*Vch*USA3) inhibit the conjugation of A/C plasmids via the fertility inhibition factor *ctiC*, a small conserved gene located downstream of *traG*. *ctiC* is broadly conserved in SGI1-like IMEs dwelling in several pathogenic species of *Enterobacteriaceae* and marine bacteria (*Vibrio*, *Shewanella*) (25, 26). For instance, *ctiC* is present in the IME GI-15 (Supplementary Fig. S2), which propagated ARGs in clinical isolates of *V. cholerae* O1 El Tor during the second epidemic wave of the 7^th^ cholera pandemic (80, 81). The presence of GI-15 and IncC plasmids in pandemic strains appears to have an inverse correlation (81), suggesting *ctiC* may play a crucial role in limiting the transmission of A/C plasmids among pathogens. CtiC is distantly related to and has a predicted 3D structure resembling the CTD of the MOB_H12_ relaxase TraI encoded by A/C and related untyped conjugative plasmids (Supplementary Fig. S4). By mimicking the CTD of TraI, CtiC interferes with relaxosome assembly via competition for the essential mobilization factor MobI (Fig. 7), which is necessary for *oriT* recognition (8, 19, 68). To the best of our knowledge, CtiC is the only known fertility inhibition factor interfering with relaxosome assembly.

**Figure 7.**
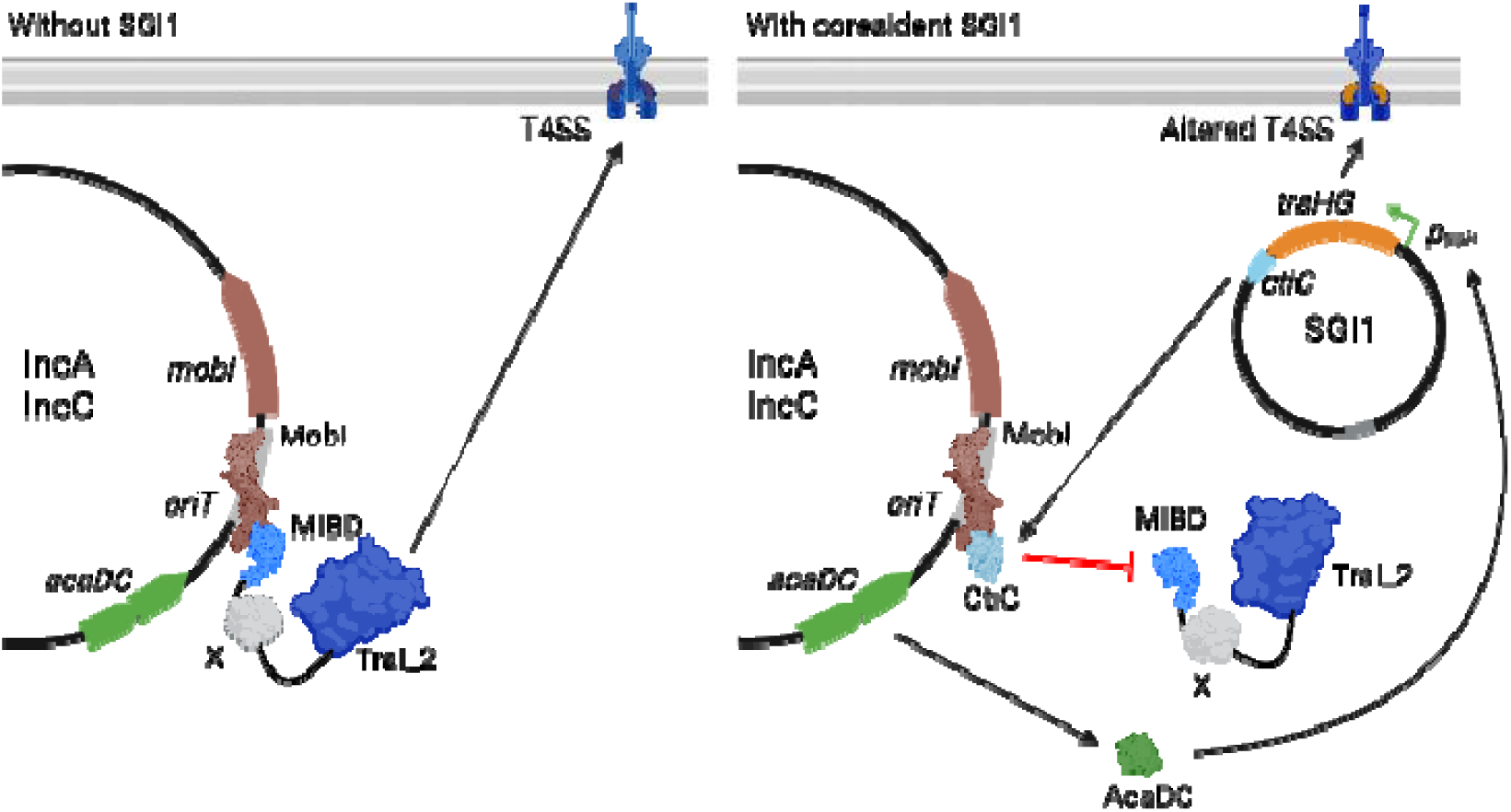
Proposed model of fertility inhibition of IncA and IncC plasmids mediated by SGI1-like IMEs. CtiC competes with MIBD of TraI to bind to MobI, preventing transfer initiation at *oriT* of A/C plasmids when SGI1 is present. SGI1 encodes its own relaxase MpsAB, unrelated to the MOB_H12_ relaxase TraI. Hence, MpsAB is not subjected to inhibition by CtiC. X is a folded domain of unknown function.

Except for the relaxases TcpM of pCW3 and MspA of SGI1, which are related to site-specific tyrosine recombinases (DNA_BRE_C superfamily) (27, 69), known relaxases are phylogenetically classified into seven major MOB families (70, 71). MOB_H_ relaxases are the only ones that harbour an N-terminal relaxase domain (HD hydrolase TraI_2), yet their mechanism of action remains poorly characterized. The MOB_H_ family is subdivided into two major clades (70). The MOB_H2_ clade contains relaxases encoded by ICE*clc* of *Pseudomonas putida* and the Gonococcal Genomic Island (GGI) of *Neisseria gonorrhoeae* (70). MOB_H2_ relaxases contain a TraI_2_C CTD of unknown function that forms a globular structure (72). The MOB_H1_ clade is subdivided into two sub-clades. MOB_H11_ includes relaxases encoded by IncHI1 and IncHI2 plasmids. MOB_H12_ contains the relaxases of A/C, IncT, and IncP-7 plasmids and SXT/R391 elements (70). Before this work, no conserved functional domain, except for the catalytic TraI_2 domain, was identified in MOB_H1_ relaxases. Here, we identified the MobI-binding domain (MIBD) at the C-terminus of TraI, a domain encoded within many diverse plasmids found in a broad range of bacterial species (Supplemental Fig. S3). By binding to the ancillary mobilization factor Mob, MIBD provides selectivity for the recognition and processing of *oriT*. Indeed, the sole replacement of MIBD in TraI of pVCR94 allowed us to change its selectivity towards MobI and *oriT* of pAQU1 (Fig. 5E). The CTDs of MOB_H1_ and MOB_H2_ relaxases are strikingly different (Supplementary Fig. S5), and MOB_H2_ relaxases do not seem to require a MobI homolog to initiate DNA transfer. The CTDs of MOB_H12_ and MOB_H11_ relaxases are also dissimilar; however, both require a MobI-like mobilization factor (e.g., TraH of R27 (74)), suggesting they play a similar role.

Genes coding for distant homologues of MobI have been found in IMEs unrelated to SGI1 (e.g., MGI*Vch*Hai6) (23). These IMEs lack a relaxase gene but are mobilized by IncC plasmids in a TraI-dependent fashion. All have an *oriT* that is unrelated to that of IncC plasmids and lies upstream of *mobI* (73). Hence, MobI proteins presumably play an integral part in the specific recognition of *oriT*s by TraI. Moreover, MIBD exhibits some versatility, being able to accommodate MobI adaptor proteins that diverge from its cognate MobI. We showed that MobI interacts with itself to form a multimeric complex (Fig. 4). Based on this observation, we constructed using Alphafold a dimeric MobI complex, a likely conformation for a DNA-binding protein, that we associated with two subunits of TraI (MIBD domain), two subunits of CtiC, or a combination of both (Fig. 8). Metrics aimed at evaluating interchain interactions in predicted structures (ipSAE>0.6 and ipTM>0.6) suggest stronger interactions between MobI and CtiC than between MobI and MIBD, which correlates with our BACTH assays (Fig. 4 and Supplemental Table S3). The mutant pVCR94 *traI*Δ*MIBD*::*ctiC* transferred poorly in our assays, suggesting that residues that increase the interactions between MobI and TraI could be detrimental to conjugative transfer (Fig. 5D). We predict that, like the MobI_2_-CtiC_2_ complex, the TraI-MobI_2_-CtiC complex would be ineffective if, like for the F plasmid (MOB_F_ family of relaxases), two subunits of TraI are required for DNA processing. Further biochemical characterization will be necessary to confirm this model, and is ongoing.

**Figure 8.**
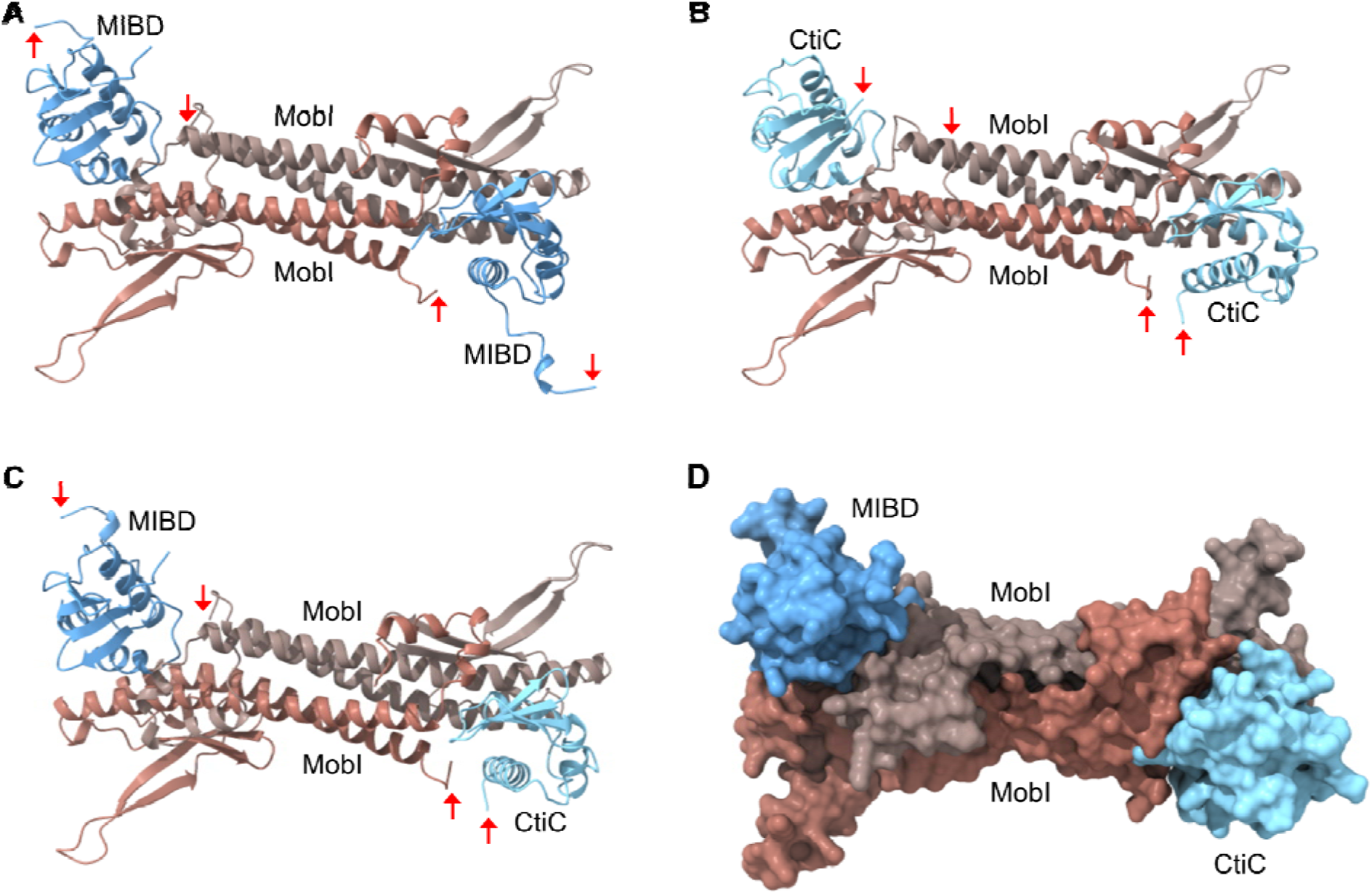
Predicted quaternary complexes formed between MobI, MIBD, and CtiC. Alphafold2 predictions of MobI_2_-MIBD_2_ (**A**), MobI_2_-CtiC_2_ (**B**), and MIBD-MobI_2_-CtiC (**C**) complexes. The red arrows indicate the N-terminus of chains. (**D**) Molecular surface prediction of an MIBD-MobI_2_-CtiC complex. Metrics of interchain interactions are presented in Supplemental Table S3.

We found that *ctiC* of SGI1 is part of the *traHG* operon driven by the *P_traH_* promoter, which is part of the AcaCD regulon, confirming previous reports (14, 30). This regulation is conserved in IME*Vch*USA3 and presumably all distantly related IMEs. Hence, *ctiC* genes are expressed in conditions that trigger the expression of the transfer genes of A/C plasmids, to which belongs the relaxase gene *traI* (14). *P_traH_* also reacts to SgaCD, the SGI1-encoded homologues of AcaCD, whose expression is activated by the SOS response (60, 75). Activation of *ctiC* expression by SgaCD could explain the slight enrichment of mini-Tn*5* insertions in *sgaDC* in our TraDIS experiment (Fig. 1B). The expression levels of *ctiC* and *traI* are strikingly similar in inducing conditions; however, *ctiC* and *ctiC3* are constitutively expressed at low levels, even in the absence of a helper plasmid (Fig. 6B). This observation suggests that SGI1-like IMEs prime the cells to quench the conjugative transfer of their helper plasmid as soon as it enters the cell.

The importance of the region encompassing *traN* to *traHG* in the lifecycle of SGI1-like IMEs has long been a matter of controversy (13, 29). Others pointed out that SGI1 variants lacking this 9.7-kb region, such as SGI1-KL1, remain mobilizable despite the absence of three T4SS subunit-encoding genes, the gene S010, now known as *ctiC*, and the SOS-response-controlled transcriptional activator genes *sgaDC* (76, 77). Although this observation is accurate in laboratory conditions, it underestimates the impact of *ctiC* on the evolutionary success of SGI1-related IMEs in a natural context. Indeed, our results using the *tra_S_* and *traHG* mutants of SGI1 undeniably confirm that the loss of this region drastically impairs SGI1 mobilization (Fig. 2C). *traHG* replacement accounts for the enhanced conjugation phenotype, but the precise mechanism remains unclear and is currently under investigation. *ctiC* prevents the helper plasmid from benefiting from the improved conjugation efficiency (Fig. 2C). By reducing the odds of cotransfer, CtiC allows the IME to stabilize in its new host. As long as the helper plasmid coexists in the cell, the activator AcaCD elicits the expression of the excisionase of the IME, which remains excised and is hence prone to loss despite its ability to replicate (78, 79). Like SgaCD (60), TraH, TraG, and CtiC are essential components of the SGI1 lifecycle. The weaker impact of *ctiC3* deletion in IME*Vch*USA3 on the helper plasmid compared to *ctiC* deletion in SGI1 (compare Fig. 1D and 3D) likely stems from the absence of *traH* and *sgaD* in IME*Vch*USA3 (26).

Fertility inhibition is another parasitic behaviour exhibited by SGI1-like IMEs towards A/C plasmids. SGI1 TraG-mediated reshaping of the T4SS allows SGI1 to dodge IncC (and IncA) entry exclusion mediated by *eexC* (13, 20). Entry exclusion prevents the redundant transfer of a conjugative element into a recipient cell containing the same or a similar element. Entry exclusion avoidance presumably helps SGI1 to propagate efficiently between IncC^+^ or IncA^+^ cells. SGI1 also seems to exclude A/C plasmids by an unknown mechanism (30). Furthermore, SGI1 excises and replicates in the presence of a helper plasmid while destabilizing it (78, 79). Murányi *et al.* (82) showed that SGI1 co-opts the ParAB partitioning system of IncC plasmids to ensure the proper segregation of its replicative form in daughter cells, eliciting helper IncC plasmid loss (78). By directly inhibiting IncC and IncA plasmid cotransfer, *ctiC* contributes to the antagonistic arsenal used by SGI1-like IMEs to curb the spread of their helper plasmids on multiple fronts. By highlighting the significance of MIBD in the TraI-mediated processing of *oriT* through its interaction with MobI, CtiC paves the way for the design of conjugation inhibitors.

## Data availability

Raw sequencing data of TraDIS experiments were submitted to Genbank Sequence Read Archive (SRA) under BioProject accession number PRJNA857050 with the following BioSample accession numbers: SAMN29603706 and SAMN29603707 for *E. coli* KH95 bearing pVCR94^Km^ Δ*acr2* and SGI1^Cm^ as the donor strain (input); SAMN29603708 and SAMN29603709 for transconjugants with *E. coli* GG56 as the recipient strain (output). Complete data from aligned reads can also be visualized using the UCSC genome browser at https://genome.ucsc.edu/cgi-bin/hgHubConnect?hgHub_do_redirect=on&hgHubConnect.remakeTrackHub=on&hgHub_do_firstDb=1&hubUrl=https://g-f2b62d.6d81c.5898.data.globus.org/Burrus/Deschenes_2023_SGI1Cm/SGI1Cm.hub.txt (SGI1^Cm^) and https://genome.ucsc.edu/cgi-bin/hgHubConnect?hgHub_do_redirect=on&hgHubConnect.remakeTrackHub=on&hgHub_do_firstDb=1&hubUrl=https://g-f2b62d.6d81c.5898.data.globus.org/Burrus/Deschenes_2023_pVCR94dX3dacr2/pVCR94dX3dacr2.hub.txt (pVCR94^Km^ Δ*acr2*).

## Supplementary Data statement

Supplementary Data are available at *NAR* Online

## Acknowledgments

We are grateful to Christian Baron for the kind gift of *E. coli* BTH101 and the BACTH plasmids pUT18C and pKT25. We thank Benoît Doublet and Satoru Suzuki for the kind gifts of the IncA conjugative plasmid pRA1 and untyped conjugative plasmid pAQU1, respectively. We also thank Frédérik Leroux for technical assistance.

## References

1. Partridge, S.R., Kwong, S.M., Firth, N. and Jensen, S.O. (2018) Mobile Genetic Elements Associated with Antimicrobial Resistance. Clin Microbiol Rev, 31, e00088–17.

2. Botelho, J. and Schulenburg, H. (2021) The Role of Integrative and Conjugative Elements in Antibiotic Resistance Evolution. Trends Microbiol., 29, 8–18.

3. Castañeda-Barba, S., Top, E.M. and Stalder, T. (2024) Plasmids, a molecular cornerstone of antimicrobial resistance in the One Health era. Nat Rev Microbiol, 22, 18–32.

4. Cabezón, E., Ripoll-Rozada, J., Peña, A., Cruz, F. de la and Arechaga, I. (2015) Towards an integrated model of bacterial conjugation. FEMS Microbiology Reviews, 39, 81–95.

5. Bellanger, X., Payot, S., Leblond-Bourget, N. and Guédon, G. (2014) Conjugative and mobilizable genomic islands in bacteria: evolution and diversity. FEMS Microbiology Reviews, 38, 720–760.

6. Ares-Arroyo, M., Coluzzi, C., Moura de Sousa, J.A. and Rocha, E.P.C. (2024) Hijackers, hitchhikers, or co-drivers? The mysteries of mobilizable genetic elements. PLoS Biol, 22, e3002796.

7. Harmer, C.J. and Hall, R.M. (2015) The A to Z of A/C plasmids. Plasmid, 80, 63–82.

8. Carraro, N., Sauvé, M., Matteau, D., Lauzon, G., Rodrigue, S. and Burrus, V. (2014) Development of pVCR94ΔX from *Vibrio cholerae*, a prototype for studying multidrug resistant IncA/C conjugative plasmids. Front Microbiol, 5, 44.

9. Fricke, W.F., Welch, T.J., McDermott, P.F., Mammel, M.K., LeClerc, J.E., White, D.G., Cebula, T.A. and Ravel, J. (2009) Comparative genomics of the IncA/C multidrug resistance plasmid family. J. Bacteriol., 191, 4750–4757.

10. Ma, L., Yin, Z., Zhang, D., Zhan, Z., Wang, Q., Duan, X., Gao, H., Liang, Q., Zhao, Y., Feng, J., et al. (2017) Comparative genomics of type 1 IncC plasmids from China. Future Microbiol, 12, 1511–1522.

11. Fernández-Alarcón, C., Singer, R.S. and Johnson, T.J. (2011) Comparative genomics of multidrug resistance-encoding IncA/C plasmids from commensal and pathogenic *Escherichia coli* from multiple animal sources. PLoS One, 6, e23415.

12. Hazen, T.H., Zhao, L., Boutin, M.A., Stancil, A., Robinson, G., Harris, A.D., Rasko, D.A. and Johnson, J.K. (2014) Comparative genomics of an IncA/C multidrug resistance plasmid from *Escherichia coli* and *Klebsiella* isolates from intensive care unit patients and the utility of whole-genome sequencing in health care settings. Antimicrob Agents Chemother, 58, 4814–4825.

13. Carraro, N., Durand, R., Rivard, N., Anquetil, C., Barrette, C., Humbert, M. and Burrus, V. (2017) *Salmonella* genomic island 1 (SGI1) reshapes the mating apparatus of IncC conjugative plasmids to promote self-propagation. PLOS Genetics, 13, e1006705.

14. Carraro, N., Matteau, D., Luo, P., Rodrigue, S. and Burrus, V. (2014) The master activator of IncA/C conjugative plasmids stimulates genomic islands and multidrug resistance dissemination. PLoS Genet, 10, e1004714.

15. Roy, D., Huguet, K.T., Grenier, F. and Burrus, V. (2020) IncC conjugative plasmids and SXT/R391 elements repair double-strand breaks caused by CRISPR-Cas during conjugation. Nucleic Acids Res., 48, 8815–8827.

16. Hancock, S.J., Phan, M.-D., Luo, Z., Lo, A.W., Peters, K.M., Nhu, N.T.K., Forde, B.M., Whitfield, J., Yang, J., Strugnell, R.A., et al. (2020) Comprehensive analysis of IncC plasmid conjugation identifies a crucial role for the transcriptional regulator AcaB. Nat Microbiol, 5, 1340–1348.

17. Hancock, S.J., Phan, M.-D., Peters, K.M., Forde, B.M., Chong, T.M., Yin, W.-F., Chan, K.-G., Paterson, D.L., Walsh, T.R., Beatson, S.A., et al. (2017) Identification of IncA/C Plasmid Replication and Maintenance Genes and Development of a Plasmid Multilocus Sequence Typing Scheme. Antimicrob. Agents Chemother., 61, e01740–16.

18. Hancock, S.J., Phan, M., Roberts, L.W., Vu, T.N.M., Harris, P.N.A., Beatson, S.A. and Schembri, M.A. (2021) Characterization of DtrJ as an IncC plasmid conjugative DNA transfer component. Mol Microbiol, 116, 154–167.

19. Hegyi, A., Szabó, M., Olasz, F. and Kiss, J. (2017) Identification of *oriT* and a recombination hot spot in the IncA/C plasmid backbone. Sci Rep, 7, 10595.

20. Humbert, M., Huguet, K.T., Coulombe, F. and Burrus, V. (2019) Entry Exclusion of Conjugative Plasmids of the IncA, IncC, and Related Untyped Incompatibility Groups. J. Bacteriol., 201, e00731–18.

21. Rivard, N., Humbert, M., Huguet, K.T., Fauconnier, A., Bucio, C.P., Quirion, E. and Burrus, V. (2024) Surface exclusion of IncC conjugative plasmids and their relatives. PLoS Genet, 20, e1011442.

22. Douard, G., Praud, K., Cloeckaert, A. and Doublet, B. (2010) The *Salmonella* genomic island 1 is specifically mobilized in trans by the IncA/C multidrug resistance plasmid family. PLoS ONE, 5, e15302.

23. Carraro, N., Rivard, N., Ceccarelli, D. and Burrus, V. (2016) IncA/C Conjugative Plasmids Mobilize a New Family of Multidrug Resistance Islands in Clinical *Vibrio cholerae* Non-O1/Non-O139 Isolates from Haiti. mBio, 7, e00509–16.

24. Boyd, D., Peters, G.A., Cloeckaert, A., Boumedine, K.S., Chaslus-Dancla, E., Imberechts, H. and Mulvey, M.R. (2001) Complete nucleotide sequence of a 43-kilobase genomic island associated with the multidrug resistance region of *Salmonella enterica* serovar Typhimurium DT104 and its identification in phage type DT120 and serovar Agona. J. Bacteriol., 183, 5725–5732.

25. de Curraize, C., Siebor, E. and Neuwirth, C. (2021) Genomic islands related to *Salmonella* genomic island 1; integrative mobilisable elements in *trmE* mobilised in trans by A/C plasmids. Plasmid, 114, 102565.

26. Durand, R., Deschênes, F. and Burrus, V. (2021) Genomic islands targeting *dusA* in *Vibrio* species are distantly related to *Salmonella* Genomic Island 1 and mobilizable by IncC conjugative plasmids. PLoS Genet, 17, e1009669.

27. Kiss, J., Szabó, M., Hegyi, A., Douard, G., Praud, K., Nagy, I., Olasz, F., Cloeckaert, A. and Doublet, B. (2019) Identification and Characterization of *oriT* and Two Mobilization Genes Required for Conjugative Transfer of *Salmonella* Genomic Island 1. Front Microbiol, 10, 457.

28. Harmer, C.J., Hamidian, M., Ambrose, S.J. and Hall, R.M. (2016) Destabilization of IncA and IncC plasmids by SGI1 and SGI2 type *Salmonella* genomic islands. Plasmid, 87–88, 51–57.

29. Ambrose, S.J. and Hall, R.M. (2022) Can SGI1 family integrative mobilizable elements overcome entry exclusion exerted by IncA and IncC plasmids on IncC plasmids? Plasmid, 123–124, 102654.

30. Ambrose, S.J. and Hall, R.M. (2025) SGI1 excludes IncA and IncC plasmids. Plasmid, 133, 102743.

31. Getino, M. and de la Cruz, F. (2018) Natural and Artificial Strategies To Control the Conjugative Transmission of Plasmids. Microbiol Spectr, 6, 6.1.03.

32. Dower, W.J., Miller, J.F. and Ragsdale, C.W. (1988) High efficiency transformation of *E. coli* by high voltage electroporation. Nucleic Acids Res., 16, 6127–6145.

33. Datsenko, K.A. and Wanner, B.L. (2000) One-step inactivation of chromosomal genes in *Escherichia coli* K-12 using PCR products. Proc Natl Acad Sci U S A, 97, 6640–6645.

34. Karimova, G., Dautin, N. and Ladant, D. (2005) Interaction Network among *Escherichia coli* Membrane Proteins Involved in Cell Division as Revealed by Bacterial Two-Hybrid Analysis. J Bacteriol, 187, 2233–2243.

35. Miller, J.H. (1992) A Short Course in Bacterial Genetics: A Laboratory Manual and Handbook for Escherichia coli and Related Bacteria Cold Spring Harbor Laboratory Press, Cold Spring Harbor.

36. Remmert, M., Biegert, A., Hauser, A. and Söding, J. (2012) HHblits: lightning-fast iterative protein sequence searching by HMM-HMM alignment. Nat Methods, 9, 173–175.

37. Mirdita, M., Steinegger, M. and S”oding, J. (2019) MMseqs2 desktop and local web server app for fast, interactive sequence searches. Bioinformatics, 35, 2856–2858.

38. Lemoine, F., Correia, D., Lefort, V., Doppelt-Azeroual, O., Mareuil, F., Cohen-Boulakia, S. and Gascuel, O. (2019) NGPhylogeny.fr: new generation phylogenetic services for non-specialists. Nucleic Acids Research, 47, W260–W265.

39. Capella-Gutiérrez, S., Silla-Martínez, J.M. and Gabaldón, T. (2009) trimAl: a tool for automated alignment trimming in large-scale phylogenetic analyses. Bioinformatics, 25, 1972–1973.

40. Edgar, R.C. (2004) MUSCLE: multiple sequence alignment with high accuracy and high throughput. Nucleic Acids Res., 32, 1792–1797.

41. Guindon, S., Dufayard, J.-F., Lefort, V., Anisimova, M., Hordijk, W. and Gascuel, O. (2010) New algorithms and methods to estimate maximum-likelihood phylogenies: assessing the performance of PhyML 3.0. Syst. Biol., 59, 307–321.

42. Lefort, V., Longueville, J.-E. and Gascuel, O. (2017) SMS: Smart Model Selection in PhyML. Molecular Biology and Evolution, 34, 2422–2424.

43. Lemoine, F., Domelevo Entfellner, J.-B., Wilkinson, E., Correia, D., Dávila Felipe, M., De Oliveira, T. and Gascuel, O. (2018) Renewing Felsenstein’s phylogenetic bootstrap in the era of big data. Nature, 556, 452–456.

44. Letunic, I. and Bork, P. (2024) Interactive Tree of Life (iTOL) v6: recent updates to the phylogenetic tree display and annotation tool. Nucleic Acids Research, 52, W78–W82.

45. Mirdita, M., Schütze, K., Moriwaki, Y., Heo, L., Ovchinnikov, S. and Steinegger, M. (2022) ColabFold: making protein folding accessible to all. Nat Methods, 19, 679–682.

46. Goddard, T.D., Huang, C.C., Meng, E.C., Pettersen, E.F., Couch, G.S., Morris, J.H. and Ferrin, T.E. (2018) UCSF ChimeraX: Meeting modern challenges in visualization and analysis: UCSF ChimeraX Visualization System. Protein Science, 27, 14–25.

47. Krissinel, E. and Henrick, K. (2004) Secondary-structure matching (SSM), a new tool for fast protein structure alignment in three dimensions. Acta Crystallogr D Biol Crystallogr, 60, 2256–2268.

48. Holm, L. (2022) Dali server: structural unification of protein families. Nucleic Acids Research, 50, W210–W215.

49. Robert, X. and Gouet, P. (2014) Deciphering key features in protein structures with the new ENDscript server. Nucleic Acids Research, 42, W320–W324.

50. Dunbrack, R.L. (2025) Rēs ipSAE loquunturLJ: What’s wrong with AlphaFold’s ipTM score and how to fix it.10.1101/2025.02.10.637595.

51. R Core Team (2022) R: A Language and Environment for Statistical Computing R Foundation for Statistical Computing, Vienna, Austria.

52. Hothorn, T., Bretz, F. and Westfall, P. (2008) Simultaneous Inference in General Parametric Models. Biom. J., 50, 346–363.

53. Wickham, H. (2016) ggplot2: Elegant Graphics for Data Analysis Springer-Verlag New York.

54. Kolde, R. (2018) pheatmap: Pretty Heatmaps.

55. Cabezón, E., Sastre, J.I. and de la Cruz, F. (1997) Genetic evidence of a coupling role for the TraG protein family in bacterial conjugation. Mol. Gen. Genet., 254, 400–406.

56. Núñez, B. and De La Cruz, F. (2001) Two atypical mobilization proteins are involved in plasmid CloDF13 relaxation. Mol. Microbiol., 39, 1088–1099.

57. Yan, J., Beattie, T.R., Rojas, A.L., Schermerhorn, K., Gristwood, T., Trinidad, J.C., Albers, S.V., Roversi, P., Gardner, A.F., Abrescia, N.G.A., et al. (2017) Identification and characterization of a heterotrimeric archaeal DNA polymerase holoenzyme. Nat Commun, 8, 15075.

58. Aravind, L., Anantharaman, V., Balaji, S., Babu, M. and Iyer, L. (2005) The many faces of the helix-turn-helix domain: Transcription regulation and beyond. FEMS Microbiology Reviews, 29, 231–262.

59. Gajiwala, K.S. and Burley, S.K. (2000) Winged helix proteins. Current Opinion in Structural Biology, 10, 110–116.

60. Durand, R., Huguet, K.T., Rivard, N., Carraro, N., Rodrigue, S. and Burrus, V. (2021) Crucial role of *Salmonella* genomic island 1 master activator in the parasitism of IncC plasmids. Nucleic Acids Res, 49, 7807–7824.

61. Koraimann, G., Teferle, K., Markolin, G., Woger, W. and Högenauer, G. (1996) The FinOP repressor system of plasmid R1: analysis of the antisense RNA control of *traJ* expression and conjugative DNA transfer. Molecular Microbiology, 21, 811–821.

62. Gasson, M.J. and Willetts, N.S. (1977) Further characterization of the F fertility inhibition systems of ‘unusual’ Fin+ plasmids. J Bacteriol, 131, 413–420.

63. Gaffney, D., Skurray, R., Willetts, N. and Brenner, S. (1983) Regulation of the F conjugation genes studied by hybridization and *tra*-*lacZ* fusion. Journal of Molecular Biology, 168, 103–122.

64. Gasson, M.J. and Willetts, N.S. (1975) Five control systems preventing transfer of *Escherichia coli* K-12 sex factor F. J Bacteriol, 122, 518–525.

65. Willetts, N. (1980) Interactions between the F conjugal transfer system and CloDF13::TnA plasmids. Molec. Gen. Genet., 180, 213–217.

66. Santini, J.M. and Stanisich, V.A. (1998) Both the *fipA* gene of pKM101 and the *pifC* gene of F inhibit conjugal transfer of RP1 by an effect on *traG*. J Bacteriol, 180, 4093–4101.

67. Maindola, P., Raina, R., Goyal, P., Atmakuri, K., Ojha, A., Gupta, S., Christie, P.J., Iyer, L.M., Aravind, L. and Arockiasamy, A. (2014) Multiple enzymatic activities of ParB/Srx superfamily mediate sexual conflict among conjugative plasmids. Nat Commun, 5, 5322.

68. Ceccarelli, D., Daccord, A., René, M. and Burrus, V. (2008) Identification of the origin of transfer (*oriT*) and a new gene required for mobilization of the SXT/R391 family of integrating conjugative elements. J. Bacteriol., 190, 5328–5338.

69. Wisniewski, J.A., Traore, D.A., Bannam, T.L., Lyras, D., Whisstock, J.C. and Rood, J.I. (2016) TcpM: a novel relaxase that mediates transfer of large conjugative plasmids from *Clostridium perfringens*: The novel relaxase TcpM. Molecular Microbiology, 99, 884–896.

70. Garcillán-Barcia, M.P., Francia, M.V. and de la Cruz, F. (2009) The diversity of conjugative relaxases and its application in plasmid classification. FEMS Microbiol. Rev., 33, 657–687.

71. Guzmán-Herrador, D.L. and Llosa, M. (2019) The secret life of conjugative relaxases. Plasmid, 104, 102415.

72. Heilers, J.-H., Reiners, J., Heller, E.-M., Golzer, A., Smits, S.H.J. and van der Does, C. (2019) DNA processing by the MOB_H_ family relaxase TraI encoded within the gonococcal genetic island. Nucleic Acids Res., 47, 8136–8153.

73. Rivard, N., Colwell, R.R. and Burrus, V. (2020) Antibiotic Resistance in *Vibrio cholerae*: Mechanistic Insights from IncC Plasmid-Mediated Dissemination of a Novel Family of Genomic Islands Inserted at *trmE*. mSphere, 5, e00748–20.

74. Lawley, T.D., Gilmour, M.W., Gunton, J.E., Standeven, L.J. and Taylor, D.E. (2002) Functional and mutational analysis of conjugative transfer region 1 (Tra1) from the IncHI1 plasmid R27. J. Bacteriol., 184, 2173–2180.

75. Pons, M.C., Praud, K., Da Re, S., Cloeckaert, A. and Doublet, B. (2022) Conjugative IncC Plasmid Entry Triggers the SOS Response and Promotes Effective Transfer of the Integrative Antibiotic Resistance Element SGI1. Microbiol Spectr, 10.1128/spectrum.02201-22.

76. de Curraize, C., Siebor, E., Varin, V., Neuwirth, C. and Hall, R.M. (2020) Two New SGI1-LK Variants Found in *Proteus mirabilis* and Evolution of the SGI1-HKL Group of *Salmonella* Genomic Islands. mSphere, 5, e00875–19.

77. Kiss, J., Nagy, B. and Olasz, F. (2012) Stability, entrapment and variant formation of *Salmonella* genomic island 1. PLoS ONE, 7, e32497.

78. Huguet, K.T., Rivard, N., Garneau, D., Palanee, J. and Burrus, V. (2020) Replication of the *Salmonella* Genomic Island 1 (SGI1) triggered by helper IncC conjugative plasmids promotes incompatibility and plasmid loss. PLoS Genet, 16, e1008965.

79. Szabó, M., Murányi, G. and Kiss, J. (2021) IncC helper dependent plasmid-like replication of *Salmonella* Genomic Island 1. Nucleic Acids Research, 49, 832–846.

80. Grim, C.J., Hasan, N.A., Taviani, E., Haley, B., Chun, J., Brettin, T.S., Bruce, D.C., Detter, J.C., Han, C.S., Chertkov, O., et al. (2010) Genome sequence of hybrid *Vibrio cholerae* O1 MJ-1236, B-33, and CIRS101 and comparative genomics with *V. cholerae*. J. Bacteriol., 192, 3524–3533.

81. Weill, F.-X., Domman, D., Njamkepo, E., Tarr, C., Rauzier, J., Fawal, N., Keddy, K.H., Salje, H., Moore, S., Mukhopadhyay, A.K., et al. (2017) Genomic history of the seventh pandemic of cholera in Africa. Science, 358, 785–789.

82. Murányi, G., Szabó, M., Acsai, K. and Kiss, J. (2024) Two birds with one stone: SGI1 can stabilize itself and expel the IncC helper by hijacking the plasmid *parABS* system. Nucleic Acids Research, 10.1093/nar/gkae050.

83. Ferrières, L., Hémery, G., Nham, T., Guérout, A.-M., Mazel, D., Beloin, C. and Ghigo, J.-M. (2010) Silent mischief: bacteriophage Mu insertions contaminate products of *Escherichia coli* random mutagenesis performed using suicidal transposon delivery plasmids mobilized by broad-host-range RP4 conjugative machinery. J. Bacteriol., 192, 6418–6427.

84. Grenier, F., Matteau, D., Baby, V. and Rodrigue, S. (2014) Complete genome sequence of *Escherichia coli* BW25113. Genome Announc, 2, pii: e01038-14.

85. Singer, M., Baker, T.A., Schnitzler, G., Deischel, S.M., Goel, M., Dove, W., Jaacks, K.J., Grossman, A.D., Erickson, J.W. and Gross, C.A. (1989) A collection of strains containing genetically linked alternating antibiotic resistance elements for genetic mapping of *Escherichia coli*. Microbiol. Rev., 53, 1–24.

86. Garriss, G., Waldor, M.K. and Burrus, V. (2009) Mobile antibiotic resistance encoding elements promote their own diversity. PLoS Genet., 5, e1000775.

87. Datta, S., Costantino, N. and Court, D.L. (2006) A set of recombineering plasmids for gram-negative bacteria. Gene, 379, 109–115.

88. Cherepanov, P.P. and Wackernagel, W. (1995) Gene disruption in *Escherichia coli*: TcR and KmR cassettes with the option of Flp-catalyzed excision of the antibiotic-resistance determinant. Gene, 158, 9–14.

89. Aoki, T., Egusa, S., Ogata, Y. and Watanabe, T. (1971) Detection of Resistance Factors in Fish Pathogen *Aeromonas liquefaciens*. Journal of General Microbiology, 65, 343–349.

90. Nonaka, L., Maruyama, F., Miyamoto, M., Miyakoshi, M., Kurokawa, K. and Masuda, M. (2012) Novel conjugative transferable multiple drug resistance plasmid pAQU1 from *Photobacterium damselae* subsp. *damselae* isolated from marine aquaculture environment. Microbes Environ., 27, 263–272.

91. Guzman, L.M., Belin, D., Carson, M.J. and Beckwith, J. (1995) Tight regulation, modulation, and high-level expression by vectors containing the arabinose *P_BAD_* promoter. J. Bacteriol., 177, 4121–4130.

92. Karimova, G., Ullmann, A. and Ladant, D. (2001) Protein-protein interaction between *Bacillus stearothermophilus* tyrosyl-tRNA synthetase subdomains revealed by a bacterial two-hybrid system. J Mol Microbiol Biotechnol, 3, 73–82.

