## Supplemental Text S1 for "Conjugative transfer inhibition of IncA and IncC plasmids by pervasive SGI1-like elements via relaxosome assembly interference"

Running Head: Fertility inhibition of IncA and IncC plasmid by SGI1

*^1^Present address: Centre de recherche du CHU de Québec-Université Laval, Québec, Canada

*^2^Present address: CNRS, LCPME, UMR 7564, Université de Lorraine, Villers-lès-Nancy, France

*^3^Present address: Laboratoire d'Ingénierie des Systèmes Macromoléculaires (LISM), Aix-Marseille Université, CNRS, UMR 7255, Marseille, France

**Supplemental Text 1**

### Plasmid and strain constructions

Plasmids and oligonucleotides used in this study are listed in Table 1 and Supplementary Table S9. The Δ*ctiC* mutation was constructed in SGI1^Cm^ and SGI1^Km^ using primer pair SGI1del010_vb2_r/SGI1del010_B.for and pKD4 and pKD3 as the templates, respectively. SGI1^Km^ Δ*traHG*-*ctiC* was constructed using primer pair SGI1del010_vb2_r/SGIdelS012.for and pKD3 as the template. The deletion of *ctiC3* in IME*Vch*USA3^Km^ was constructed using primer pair ctiC2_W_F/CtiC2_W_R and pKD3 as the template. *ctiC*’-‘*lacZ* fusions in SGI1^Km^ and IME*Vch*USA3^Km^ and *mobI*’-‘*lacZ* and *traI*’-‘*lacZ* fusions in pVCR94^Cm^ were constructed using primer pairs oFD15/oFD16, oFD138r/oFD138f, oCC2R/mobI-lacZF, and oCC1R/traI-lacZF, respectively, and pVI42B as the template. In these fusions, the fifth codon of *ctiC, mobI*, and *traI* was fused to the eighth codon of *lacZ*. The λRed recombination system was expressed using pSIM6 or pSIM19 (1). When appropriate, resistance cassettes were excised from the resulting constructions using the Flp-encoding plasmid pCP20 (2). All deletions were validated by antibiotic profiling and PCR.

MIBD substitutions in *traI* of pVCR94^Cm^ were constructed by λRed- and Cas9-mediated recombineering. Briefly, an *aph* (Km) resistance cassette was introduced into the coding sequence of TraI’s CTD (amino acid residues 912-981) using the λRed recombination system expressed from pSIM6, the primer pair oVB22/oVB23, and pKD4 as the template. The sgRNA-expressing plasmid pAC018ΔR::*aph* was constructed by consecutively removing an sgRNA duplication and replacing the remaining sgRNA with an *aph*-targeting sgRNA using the Q5 Site-Directed Mutagenesis Kit (New England Biolabs) with primer pairs D_sgRNAAC018.for/D_sgRNAAC018.rev and oVB5/oAC776, and pAC018 as the template, following the manufacturer’s instructions. For the ΔMIBD::*ctiC* and ΔMIBD::*unk* mutants, *ctiC* or *unk* without their start codon were amplified using primer pairs oFD113f/oFD113r or oFD226F/oFD226R and SGI1^Km^ or pUC57-*unk* as the templates, respectively. For the MIBD_pAQU1_ mutant of *traI*, MIBD_pAQU1_ was amplified using primer oVB36/oVB37 and pAQU1 as the template. The substitutions of MIBD::*aph* by *ctiC*, *unk*, and MIBD_pAQU1_ were then obtained by concomitant activities of λRed and Cas9 expressed from pREDCas9 (3) and the *aph*-targeting sgRNA expressed from pAC018ΔR::*aph*.

*ctiC2* of IE*Vch*USA2 and *unk* of IE*Vch*USA5 with their native Shine-Dalgarno sequences were obtained by DNA synthesis (Bio Basic) of the fragments corresponding to positions 32,034 to 32,425 and 25,854 to 26,563 of the contigs [MIPA01000024.1](https://www.ncbi.nlm.nih.gov/nuccore/MIPA01000024) (*V. cholerae* 692-79 NODE_17_ and [NMTM01000021.1](https://www.ncbi.nlm.nih.gov/nuccore/NMTM01000021) (*V. cholerae* OYP2C05), respectively, cloned into pUC57-Amp. Plasmids pBAD30-*ctiC*, pBAD30-*ctiC2*, pBAD30-*ctiC*3, and pBAD30-*unk* were constructed by PCR amplification of *ctiC* genes using primer pairs SGI1S010EcoRI.rev/SGI1S010EcoRI.for, oFD101F/oFD101R, ctiC2_F/ctiC2_R and oFD100F/oFD100R, and SGI1^Km^, pUC57-*ctiC2*, IME*Vch*USA3^Km^ and pUC57-*unk* as the templates, respectively. The amplified fragments were then digested by EcoRI and cloned into EcoRI-digested pBAD30.

The BACTH plasmids pKT25-*ctiC*, pUT18C-*ctiC,* pKT25-*mobI*, pUT18C-*mobI,* pKT25-*traI*Δ, and pUT18C-*traI*Δ were constructed by EcoRI/XbaI-cloning into pKT25 and pUT18C of PCR-amplified *ctiC*, *mobI*, and *traI*Δ coding sequences using primer pairs SGI1S010XbaIf/SGI1S010EcoRI.rev, 94MobIXbaIf/94mobIEcoRI.rev, and oFD85R/oFD85V2, respectively. The pUT18C-*traI*ΔΔ and pUT18C-*MIBD* mutants were constructed using the Q5 site-directed mutagenesis kit (New England Biolabs) according to the manufacturer’s instructions with primer pairs oFD121r/oFD127f and oFD116r/oFD117f, respectively, and pUT18C-*traI*Δ as the template.

pMOB was constructed by amplifying the *oriT*-*mobI* region of pAQU1 (186,528-187,606 of Genbank [NC_016983.1](https://www.ncbi.nlm.nih.gov/nuccore/NC_016983.1/)) using primer pair oVB32/oVB33 and cloning the resulting fragment into EcoRI-digested pACYC184Δ*cat2*. pACYC184Δ*cat2* was obtained by Q5 site-directed mutagenesis of pACYC184Δ*cat* using the primer pair oVB30/oVB31 to introduce an EcoRI restriction site.

### Transposon-directed insertion sequencing (TraDIS)

pFG036 codes for a thermosensitive cI transcriptional repressor. pFG051 is a 𝚷-dependent RP4-mobilizable plasmid carrying the Tn*5* transposase gene repressed by cI and a mini-Tn*5* (Sp) transposon. pFG051 was transferred by conjugation from MFD*pir+* to KH95 bearing pVCR94^Km^ Δ*acr2* *trmE*::SGI1^Cm^ in a 2-h mating experiment at 30°C on LB agar plates supplemented with DAP in duplicates. The mating mixture was then entirely spread onto 40 large LB agar plates (150 mm) supplemented with Rf, Km Cm, and Sp. Plates were incubated until near confluence (10k to 40k CFUs) to select clones carrying mini-Tn*5* (Sp) insertions. After overnight incubation at 37°C, Rf Km Cm Sp-resistant colonies were collected using a cell scraper and resuspended in LB broth. The collected sample, designated as the ‘input library’, was washed, then resuspended in 4.5 ml of LB broth and cryopreserved. Total DNA of a 1.5 ml aliquot of the input library was extracted for sequencing. Another 1 ml aliquot of input library was used to inoculate 50 ml of LB broth supplemented with Rf, Km Cm, and Sp, and grown overnight at 37°C. The resulting culture was used as the donor in a mating assay, mixed in equal volumes with *E. coli* GG56 (Nx^r^). After a 5-h incubation at 37°C, mating mixtures were spread onto 30 large LB agar plates (150 mm) supplemented with Nx, Km, Cm, and Sp to select for mini-Tn*5* (Sp) insertions in SGI1^Cm^ that allowed the co-transfer and establishment of both SGI1^Cm^ and pVCR94^Km^ into the recipients. After overnight incubation at 37°C, Nx Km Cm Sp-resistant colonies were collected and subsequently resuspended in LB broth, washed and resuspended in 4.5 ml of LB broth and cryopreserved. These samples were designated as 'output libraries'. Total DNA of a 1.5 ml aliquot of the output libraries was extracted and used for sequencing.

**References**

1. Datta,S., Costantino,N. and Court,D.L. (2006) A set of recombineering plasmids for gram-negative bacteria. *Gene*, **379**, 109–115.

2. Cherepanov,P.P. and Wackernagel,W. (1995) Gene disruption in *Escherichia coli*: TcR and KmR cassettes with the option of Flp-catalyzed excision of the antibiotic-resistance determinant. *Gene*, **158**, 9–14.

3. Li,Y., Lin,Z., Huang,C., Zhang,Y., Wang,Z., Tang,Y.-J., Chen,T. and Zhao,X. (2015) Metabolic engineering of *Escherichia coli* using CRISPR-Cas9 meditated genome editing. *Metab Eng*, **31**, 13–21.

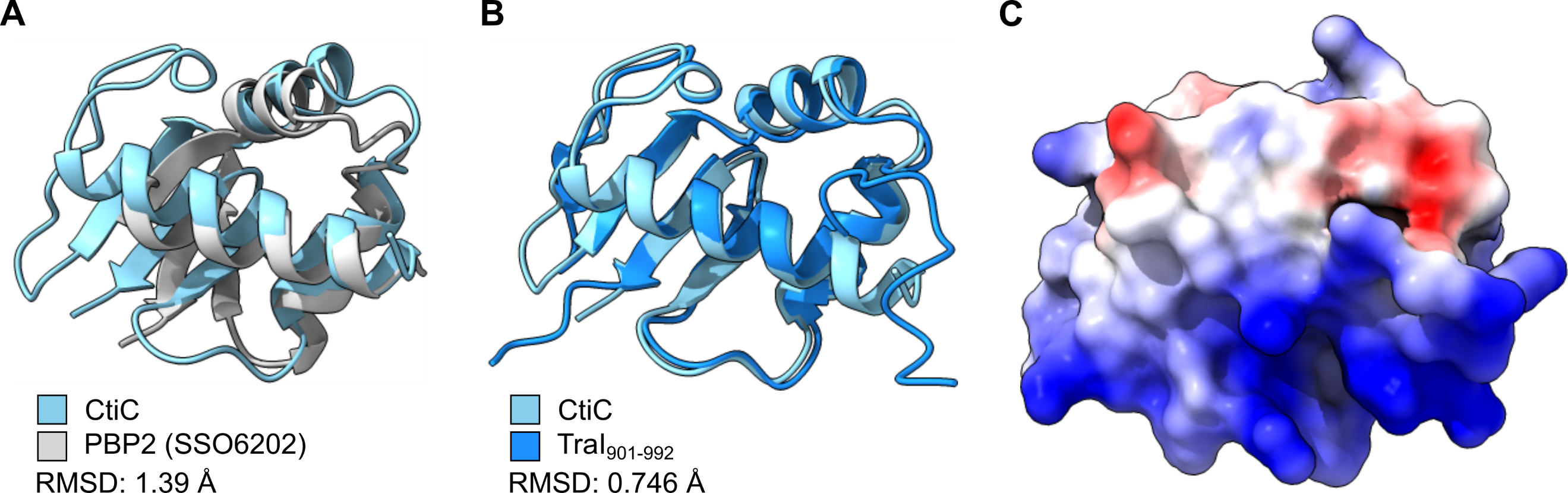

**Figure S1. CtiC exhibits a winged helix fold.** (**A**) and (**B**) Comparisons of the predicted 3D structure of CtiC with the crystal structure of PBP2 (SSO6202, PDB: 5n41), a subunit of the heterotrimeric archaeal DNA polymerase holoenzyme PolB1 from *Sulfolobus solfataricus*, or the predicted 3D structure of the 92 C-terminal sequence of TraI (TraI_901-992_) of pVCR94. Superimpositions were made using the Matchmaker tool in ChimeraX 1.9. Root mean square deviations (RMSD) between 24 (CtiC/PBP2) and 67 (CtiC/TraI_901-992_) pruned atom pairs are shown. (**C**) Predicted Coulombic electrostatic potential of CtiC. The surface was coloured from red for negative potential through white to blue for positive potential using the Coulombic command in ChimeraX 1.9.

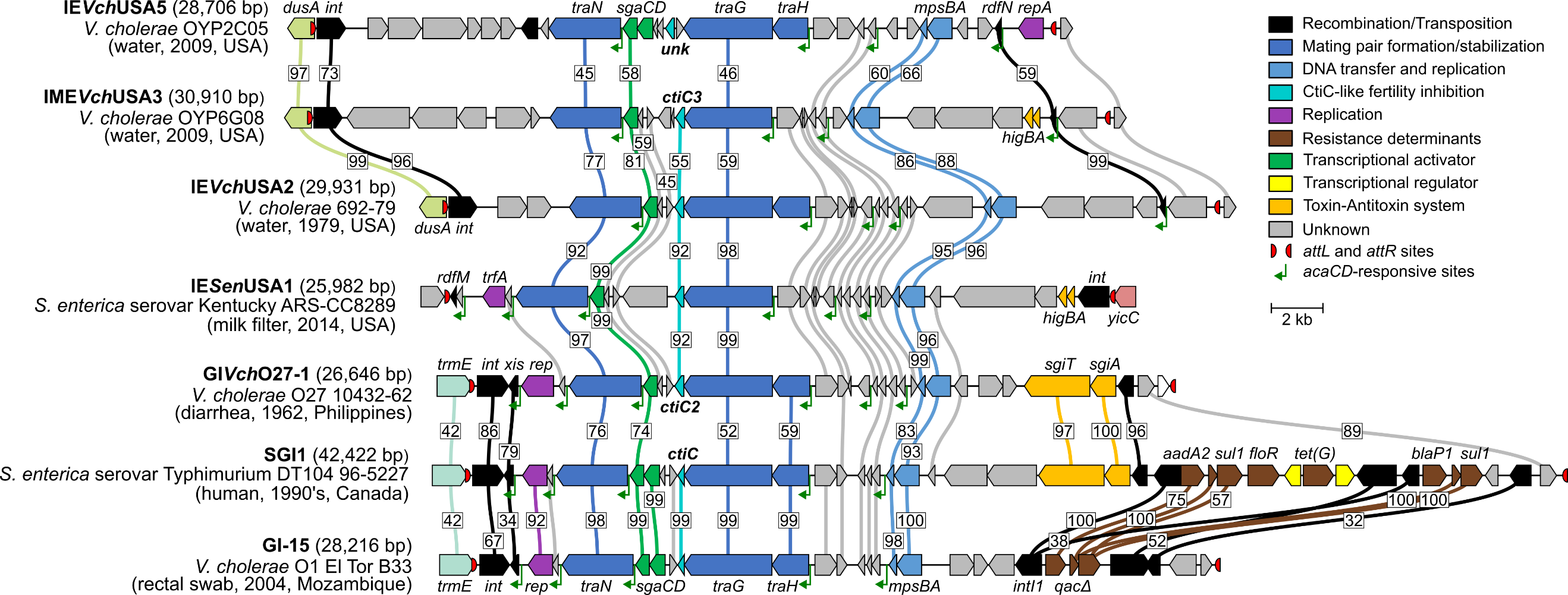

**Figure S2. Comparison of the genetic context of *ctiC*-like genes in diverse SGI1-like IMEs.** IMEs are drawn to scale. ORFs with similar functions are colour-coded as indicated in the legend. Identities between homologous proteins are shown only for genes of known function between connected ORFs. GenBank accession numbers are as follows: IE*Vch*USA5 (NZ_NMTM01000021), IME*Vch*USA3 (NZ_NMSY01000009), IE*Vch*USA2 (MIPA01000024), IE*Sen*USA1 (NZ_MCPS01000044), GI*Vch*O27-1 (CP010812), SGI1 (AF261825), and GI-15 (AAWE01000022). The respective insertion site of each IME (5’ end of *dusA*, and 3’ end of *yicC* or *trmE*) is shown.

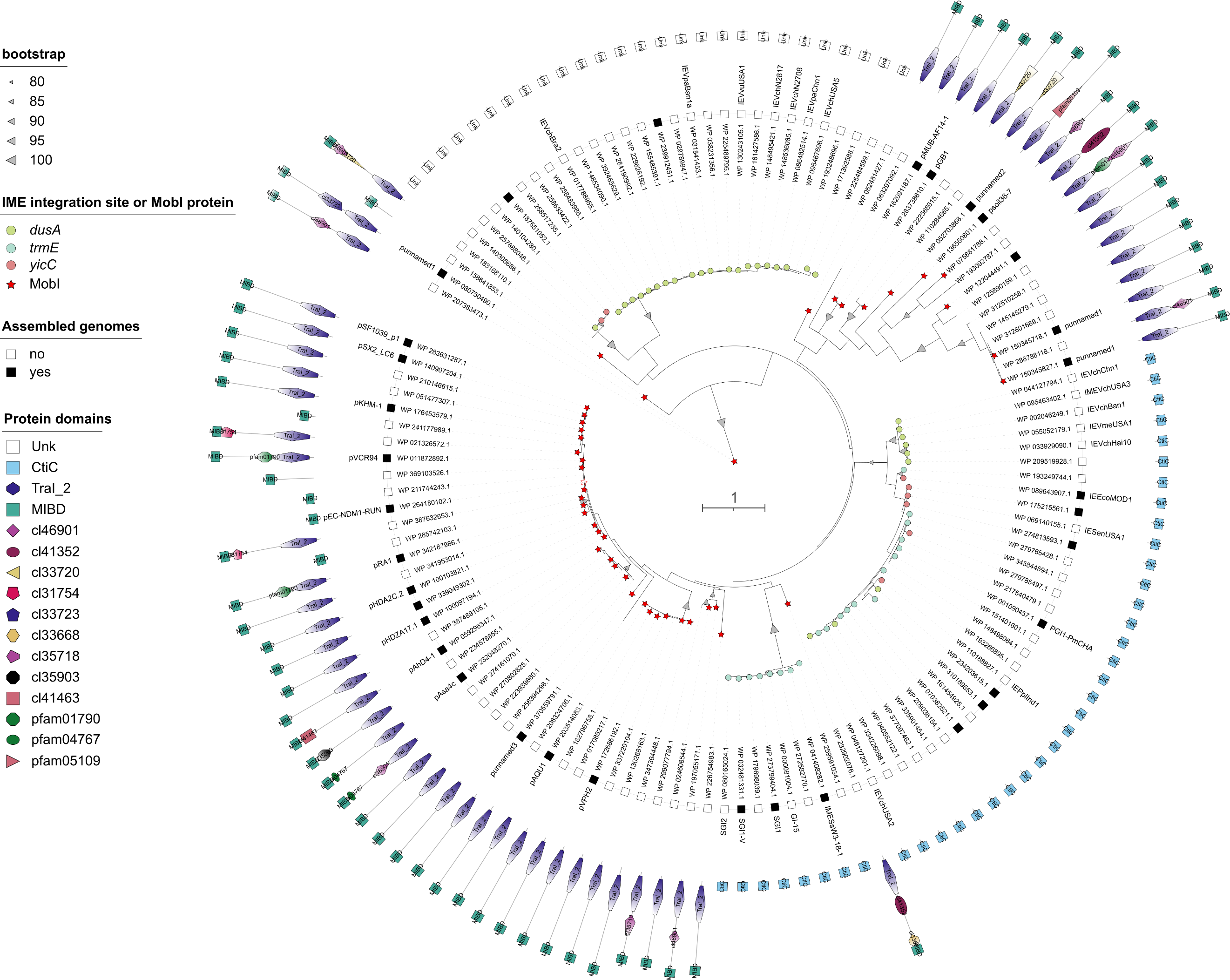

**Figure S3. Maximum likelihood phylogenetic analysis of CtiC homologues.** Bootstrap supports are indicated as grey arrowheads on the branches only when >80%. Branch lengths represent the number of substitutions per site over 80 sites. The best matrix (LG substitution model +G) was selected on the BIC criterion. Taxa or lineages corresponding to IMEs integrated at *dusA*, *trmE*, and *yicC* are shown by green, blue, and red disks, respectively. Red stars indicate lineages where a *mobI* gene is located on the same DNA molecule as the gene coding for MIBD-bearing protein. From inner to outer wheel: leaf labels shown as Genbank protein accession numbers; draft (◻) or assembled (◼) genome; the name of the plasmid or IME when known; conserved protein domains. The tree was displayed with iTOL and is available at <https://itol.embl.de/shared/4nn1aC65aoqP>. The complete description of taxa is available in Supplementary Table S2.

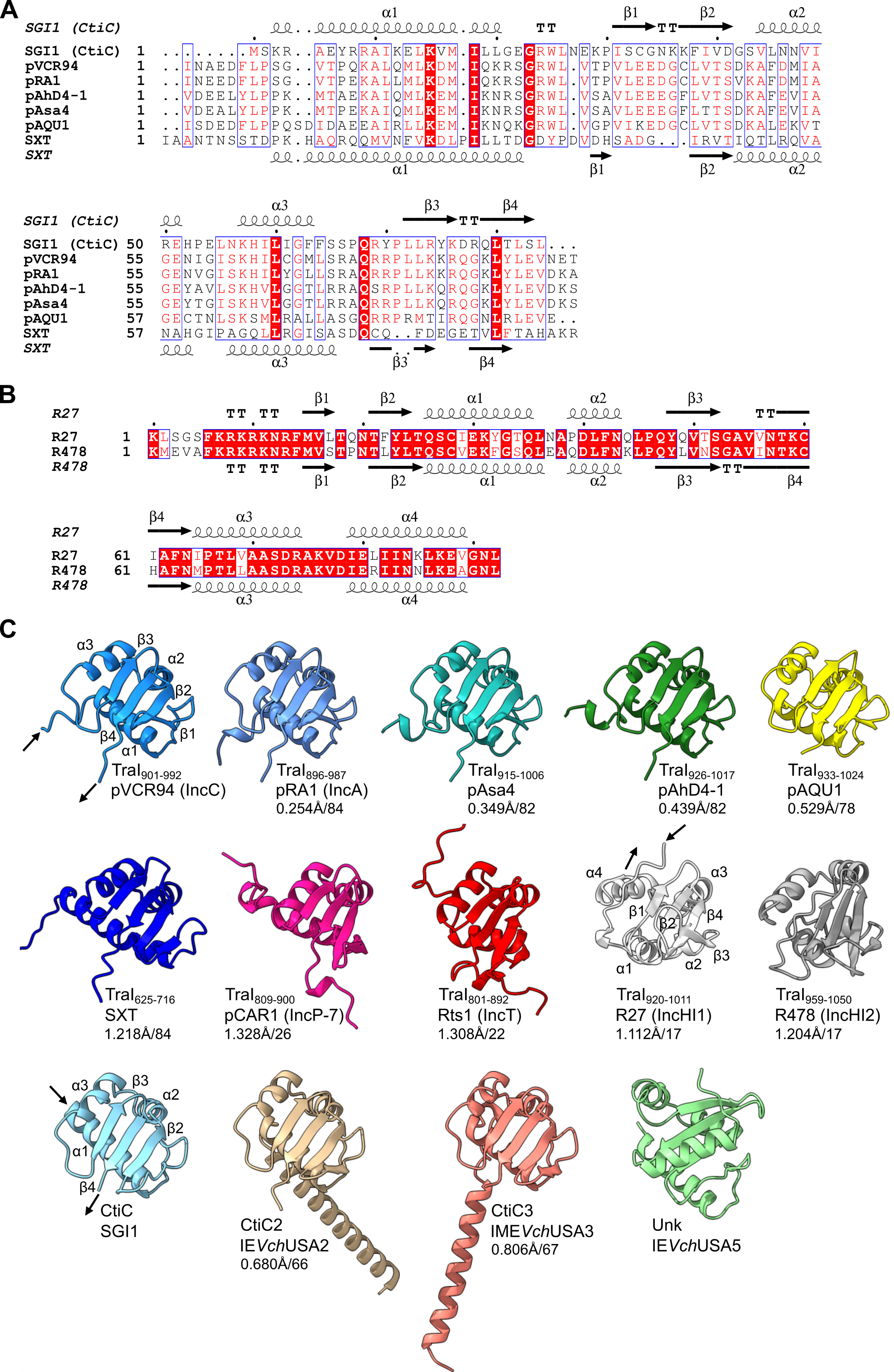

**Figure S4. Comparison of the predicted structures of MOB_H1_ relaxase CTDs and CtiC proteins.** (**A**) Muscle alignment of the primary sequences of SGI1 CtiC and the last 92 C-terminal residues of the MOB_H12_ relaxases of IncA (pRA1), IncC (pVCR94), related untyped conjugative plasmids (pAsa4, pAhD4-1, pAQU1), and SXT. The predicted secondary structures of CtiC and CTD of the relaxase encoded by SXT are depicted above and below the alignment, respectively. (**B**) Muscle alignment of the primary sequences of the last 92 C-terminal residues of the MOB_H11_ relaxases of IncHI1 (R27) and IncHI2 (R478) conjugative plasmids. The predicted secondary structures of both C-termini are depicted above and below the alignment. In A and B, similarity colouring is shown based on a percentage of equivalent residues calculated considering physicochemical properties. (**C**) Comparison of the predicted 3D structure of CtiC variants and the last 92 C-terminal residues of a diverse set of MOB_H1_ relaxases. Protein structures were predicted using AlphaFold2 and compared and displayed using ChimeraX 1.9 and its Matchmaker tool. The features annotated on CtiC, TraI_901-992_ (MIBD) of pVCR94 and TraI_920-1011_ of R27 correspond to those shown in panels A and B. RMSD values over the indicated pruned atom pairs for comparisons to TraI of pVCR94 (plasmids) or CtiC of SGI1 (IMEs) are displayed below each structure, except for Unk.

**Table S1.** Oligonucleotides used in this study

| **Primer name** | **Nucleotide sequence (5' to 3')*^a^*** | **Usage** |
| --- | --- | --- |
| SGI1del010_B.for | AGGTATTTGTCCGTAGCCAAAGGAGGCCGTTAATGAGTGTAGGCTGGAGCTGCTTC | Deletion of *ctiC* in SGI1 |
| SGI1del010_vb2_r | ACAAAGGCTCCTGAGAGCCTTTGTTTTTATTCATTACATATGAATATCCTCCTTA | Deletion of *traHG*-*ctiC* or *ctiC* in SGI1 |
| SGI1deIS012.for | GTAGGCTTTCGGGTGACACGAAACCTATTGGAGCAACAGTgtgtaggctggagctgcttc | Deletion of *traHG*-*ctiC* in SGI1 |
| ctiC2_W_F | TTTGCTGTCCAACAAGGTGGAGGGAAACATGAATAAAGTGTAGGCTGGAGCTGCTTCG | Deletion of *ctiC3* in IME*Vch*USA3 |
| ctiC2_W_R | AATGCTGTATATAAAAACAGCATCCTTATCAAGCTCACATATGAATATCCTCCTTA | Deletion of *ctiC3* in IME*Vch*USA3 |
| oFD15 | AGCGTTGCACCAATGCTCGACTGGACGGACAGACATCTGGCCGTCGTTTTACAACGTCG | *ctiC*’-*‘lacZ* fusion in SGI1 |
| oFD16 | AGGGCGTGGTGAATTTGACTACTTTTTGGTGAAAAGGCAGCATTACACGTCTTGAG | *ctiC*’-*‘lacZ* fusion in SGI1 |
| oFD138f | TCCAACAAGGTGGAGGGAAACATGAATAAAGCACTGCTGGCCGTCGTTTTACAACGTCG | *ctiC3*’-*‘lacZ* fusion in IME*Vch*USA3 |
| oFD138r | AAAATGCTGTATATAAAAACAGCATCCTTATCAAGCGTGTAGGCTGGAGCTGCTTCG | *ctiC3*’-*‘lacZ* fusion in IME*Vch*USA3 |
| oCC2R | GCCAATTCAGTGGCCGCTACAGATGCTGTCATGTTGGTGTAGGCTGGAGCTGCTTC | *mobI’*-*‘lacZ* fusion in pVCR94^Cm^ |
| mobI-lacZF | AGGAATTGGGAGGGTATTGAGGTGAGTCTACCAACACTGGCCGTCGTTTTACAACGTCG | *mobI’*-*‘lacZ* fusion in pVCR94^Cm^ |
| oCC1R | CATCTCGTAGGCGAGCGGGTCATAACTCATTGTCATGTGTAGGCTGGAGCTGCT | *traI*-*‘lacZ* fusion in pVCR94^Cm^ |
| traI-lacZF | ATAGGAATATGGTCAACCTACATGCTGAAAGCCCTTCTGGCCGTCGTTTTACAACGTCG | *traI*-*‘lacZ* fusion in pVCR94^Cm^ |
| oVB22 | TTGCCGTCTGGTGTTACGCCTCAGAAAGCACTCCAGTCACGCTGCCGCAAGCACTC | Insert *aph* (Km) in *traI* of pVCR94^Cm^ |
| oVB23 | TGTTTCATTTACCTCTAAATACAATTTTCCCTGACGTGATGGCAGGTTGGGCGTCG | Insert *aph* (Km) in *traI* of pVCR94^Cm^ |
| D_sgRNAAC018.for | CTTCCTCGCTCACTGACT | Deletion of sgDNA duplicate in pAC018 |
| D_sgRNAAC018.rev | TGACAGGTTTCCCGACTG | Deletion of sgDNA duplicate in pAC018 |
| oVB5 | TCATGGCTGATGCAATGCGGGTTTTAGAGCTAGAAATAGCAAGTTAAAATAAG | Site-directed mutagenesis in pAC018 |
| oAC776 | GAATCTATTATACAGAAAAATTTTCCTGAAAGC | Site-directed mutagenesis in pAC018 |
| oFD113f | CAGAGGTTAGGGAGTTTGAGCCGCCTAAGGCGAAAACAAACCCGAAAGACAGCAAACGAGCTGAATATAGA | Substitute MIBD at position 901 of TraI of pVCR94 by *ctiC* |
| oFD113r | TCGTAGGCGAGCGGGTCATAACTCATTGTCATGTTTCATTTACCTCTAAATTATAGACTTAACGTTAACTG | Substitute MIBD at position 901 of TraI of pVCR94 by *ctiC* |
| oFD226F | GAGTTTGAGCCGCCTAAGGCGAAAACAAACCCGAAAGACAATAAGAATAGAGAATTAGTT | Substitute MIBD at position 901 of TraI of pVCR94 by *unk* |
| oFD226R | CGGGTCATAACTCATTGTCATGTTTCATTTACCTCTAAATTATGGCCTTTTAATTTTATA | Substitute MIBD at position 901 of TraI of pVCR94 by *unk* |
| oVB36 | AGAAAGGGAAGAGCCAGAGGTTAGGGAGTTTGAGCCGCCTAAAAAGCCTCAGAAGGCGAA | Substitute MIDB of pVCR94 by MIBD of pAQU1 |
| oVB37 | CACGGCATCTCGTAGGCGAGCGGGTCATAACTCATTGTCACTCCACCTCTAACCTTAAAT | Substitute MIDB of pVCR94 by MIBD pAQU1 |
| SGI1S010EcoRI.for | NNNNNNGAATTCAAGGAGGAATAATAAATGAGCAAACGAGCTGAATA | Cloning of *ctiC* into pBAD30 |
| SGI1S010EcoRI.rev | NNNNNNGAATTCTTATAGACTTAACGTTAACTGCCTA | Cloning of *ctiC* into pBAD30, pKT25 or pUT18C |
| oFD101F | GCTGGAATTCAAGGAGGAATAATAAGTGAATACGGCACTGGTC | Cloning of *ctiC2* into pBAD30 |
| oFD101R | CATCGAATTCTTAACCTTCAACTCGAAGGATTAG | Cloning of *ctiC2* into pBAD30 |
| ctiC2_F | GCTGGAATTCAAGGAGGAATAATAAATGAATAAAGCACTGATACAA | Cloning of *ctiC3* into pBAD30 |
| ctiC2_R | CATCGAATTCAAGCTCATTTAATATTTAAGGTA | Cloning of *ctiC3* into pBAD30 |
| oFD100F | GCTGGAATTCAAGGAGGAATAATAAATGATGAATAAGAATAGAGAATTAGTTTA | Cloning of *unk* into pBAD30 |
| oFD100R | CATCGAATTCTTATGGCCTTTTAATTTTATAGTCC | Cloning of *unk* into pBAD30 |
| SGI1S010XbaIf | NTCTAGAAATGAGCAAACGAGCTGAA | Cloning of *ctiC* into pKT25 or pUT18C |
| 94MobIXbaIf | NTCTAGAGGTGAGTCTACCAACAGAGCGATG | Cloning of *mobI* into pKT25 or pUT18C |
| 94mobIEcoRI.rev | NNNNNNGAATTCTCACACCTCGTCGCTATGTGTCTT | Cloning of *mobI* into pKT25 or pUT18C |
| oFD85R | NNNNNNGAATTCTCATGTTTCATTTACCTCTAAATAC | Cloning of *traI*Δ into pKT25 or pUT18C |
| oFD85V2 | NTCTAGACATGCTGAAAGCCCTTAACAAGTTATT | Cloning of *traI*Δ into pKT25 or pUT18C |
| oFD121r | AACGAACATGTCTAGAGTCG | Site-directed mutagenesis in pUT18C-*traI*Δ – removal of 1-X-2 domains |
| oFD127f | ATCAACGCGGAAGATTTC | Site-directed mutagenesis in pUT18C-*traI*Δ – removal of 1-X-2 domains |
| oFD116r | GTCTTTCGGGTTTGTTTTC | Site-directed mutagenesis in pUT18C-*traI*Δ – removal of MIBD domain |
| oFD117f | TAACTAAGTAATATGGTGCAC | Site-directed mutagenesis in pUT18C-*traI*Δ – removal of MIBD domain |
| oVB30 | NNNNGAATTCTCTTCAAATGTAGCACCTG | Add an EcoRI site in pACYC184Δ*cat* |
| oVB31 | NNNNGAATTCTAAATTGCACTGAAATCTAGA | Add an EcoRI site in pACYC184Δ*cat* |
| oVB32 | TACAGAATTCATGTTGGTCTCCTTGGTTCA | EcoRI-cloning of *oriT*-*mobI* of pAQU1 |
| oVB33 | NNNNGAATTCTTACATCACGACATTTACCA | EcoRI-cloning of *oriT*-*mobI* of pAQU1 |
| traG_i_F2 | AGGCTTCCTGGCTGTAGGTG | Amplify the *ctiC*-*traG* junction |
| ctiC_i_R | CGGATACCGTTGAGGCGACG | Amplify the *ctiC*-*traG* junction |
| traH_i_F | GTTATTGCGGGTGGGCGTGTGA | Amplify an internal *traH* fragment |
| traH_i_R | CCCCGGTAAGGTGACGATGGGA | Amplify an internal *traH* fragment |
| ctiCRT | TAGACTTAACGTTAACTGCCTA | Reverse transcription of the *ctiC* mRNA |

*^a^* Restriction sites are underlined.

**Table S3.** Comparison of the interchain interactions in the MobI_2_-MIBD_2_ (pVCR94) and MobI_2_-CtiC_2_ complexes predicted by Alphafold 2. The metrics were calculated by the ipsae.py script with pae_cutoff = 5 et dist_cutoff = 15.

| **MobI_2_-MIBD_2_ complex** | | | | | | |
| --- | --- | --- | --- | --- | --- | --- |
| **Chain 1** | **Chain 2** | **ipSAE** | **ipTM_af** | **pDockQ2** | **n0res** | **d0res** |
| MobI (A) | MobI (B) | 0.832126 | 0.860 | 0.8494 | 169 | 4.85 Å |
| MobI (A) | MIBD(D) | 0.637558 | 0.860 | 0.8537 | 81 | 3.21 Å |
| MobI (B) | MIBD (C) | 0.636961 | 0.860 | 0.8521 | 81 | 3.21 Å |
| MobI (A) | MIBD (C) | 0.678720 | 0.860 | 0.8017 | 81 | 3.21 Å |
| MobI (B) | MIBD (D) | 0.680861 | 0.860 | 0.8012 | 81 | 3.21 Å |
| MIBD (C) | MIBD (D) | 0.000000 | 0.860 | 0.0000 | 0 | 1.00 Å |
| **MobI_2_-CtiC_2_ complex** | | | | | | |
| **Chain 1** | **Chain 2** | **ipSAE** | **ipTM_af** | **pDockQ2** | **n0res** | **d0res** |
| MobI (C) | MobI (D) | 0.832957 | 0.890 | 0.8576 | 170 | 4.86 Å |
| CtiC (A) | MobI (C) | 0.730885 | 0.890 | 0.8276 | 159 | 4.70 Å |
| CtiC (B) | MobI (D) | 0.731531 | 0.890 | 0.8282 | 158 | 4.68 Å |
| CtiC (B) | MobI (C) | 0.681606 | 0.890 | 0.5129 | 84 | 3.29 Å |
| CtiC (A) | MobI (D) | 0.679500 | 0.890 | 0.4993 | 84 | 3.29 Å |
| CtiC (A) | CtiC (B) | 0.000000 | 0.890 | 0.0000 | 0 | 1.00 Å |
